# The hyperlipidaemic drug fenofibrate significantly reduces infection by SARS-CoV-2 in cell culture models

**DOI:** 10.1101/2021.01.10.426114

**Authors:** Scott P. Davies, Courtney J. Mycroft-West, Isabel Pagani, Harriet J. Hill, Yen-Hsi Chen, Richard Karlsson, Ieva Bagdonaite, Scott E. Guimond, Zania Stamataki, Marcelo Andrade De Lima, Jeremy E. Turnbull, Zhang Yang, Elisa Vicenzi, Mark A. Skidmore, Farhat Khanim, Alan Richardson

## Abstract

The SARS-CoV-2 pandemic has caused a significant number of fatalities and worldwide disruption. To identify drugs to repurpose to treat SARS-CoV-2 infections, we established a screen to measure dimerization of ACE2, the primary receptor for the virus. This screen identified fenofibric acid, the active metabolite of fenofibrate. Fenofibric acid also destabilized the receptor binding domain (RBD) of the viral spike protein and inhibited RBD binding to ACE2 in ELISA and whole cell binding assays. Fenofibrate and fenofibric acid were tested by two independent laboratories measuring infection of cultured Vero cells using two different SARS-CoV-2 isolates. In both settings at drug concentrations which are clinically achievable, fenofibrate and fenofibric acid reduced viral infection by up to 70%. Together with its extensive history of clinical use and its relatively good safety profile, these studies identify fenofibrate as a potential therapeutic agent requiring urgent clinical evaluation to treat SARS-CoV-2 infection.

**Teaser:** The approved drug fenofibrate inhibits infection by SARS-COV-2

## Introduction

Severe acute respiratory syndrome coronavirus-2 (SARS-CoV-2) is responsible for a pandemic which has cost over 1.9 million lives worldwide so far (1-3). The emergence of new virus variants with higher transmissibility rates is seeing rapid increases in infection rates and deaths across the world. Several vaccines have undergone accelerated approval and are being rolled out worldwide (4,5). Whilst the data from clinical trials is very promising, the vaccines may not be suitable in all patient groups e.g. those with hyperimmune disorders and those using immunosuppressants, and it is presently unclear whether the current vaccines offer protection to newly emerging strains of the virus. In addition, it will take considerable time to vaccinate everyone and we are yet unsure of the strength and duration of the response. Therapies are still urgently needed to manage patients who develop severe symptoms and/or require hospitalisation.

The virus gains entry to human cells by the receptor binding domain (RBD) of the viral Spike protein binding to angiotensin converting enzyme-2 (ACE2) on human cells (6,7). Although other receptors of the virus have been identified (8,9), drugs which block virus binding to ACE2 may substantially reduce virus uptake thereby reducing/relieving symptoms in patients with an active infection or reduce transmission of the virus to uninfected individuals.

Whilst the rapid escalation of the SARS-CoV-2 epidemic leaves insufficient time to develop new drugs via traditional pipelines, drug repurposing offers an expedited and attractive alternative. Drugs which are repurposed are available for immediate clinical use and their pharmacokinetic and safety profiles are usually well described. This has already proven true, with the identification that dexamethasone reduces mortality of SARS-CoV-2 patients (10) and remdesivir decreases the time needed for patients to recover from infection (11). In both these cases, although the drugs are technically being repurposed, their use still depends on the drug’s recognized mechanism of action. It is less obvious which drugs might have a novel mechanism of action and interfere with SARS-CoV-2 binding and cellular entry mediated by ACE2. To this end, we recently developed an assay to measure the viral spike protein’s receptor binding domain (RBD) binding to ACE2 (12).

Structural studies have shown that ACE2 is a dimer and that there may be multiple spike RBDs interacting with each ACE2 dimer (13). Molecular dynamic simulations have suggested considerable flexibility in ACE2 and this might allow multiple ACE2 dimers to bind to each spike trimer (14). It therefore seems reasonable that the extent of ACE2 dimerization might affect the avidity of RBD binding. Furthermore, dimerization has been shown to affect internalization of other receptors. For example, dimerization of EGF or FGF receptors promotes their endocytosis (15,16) and different mechanisms of internalization may exist for monomeric and dimeric GH receptors (17). This led to the hypothesis that drugs that altered dimerization of ACE2 might affect viral infection. In order to test this hypothesis, we developed an assay to measure dimerization of ACE2, making use of the NanoBIT protein interaction system (18). This is based on a modified luciferase (nanoluc) which has been split into two catalytically incomplete components, LgBIT and SmBIT, that must bind together to form an active luciferase. LgBIT and SmBIT associate with low affinity but when fused to other proteins that interact with each other, co-localization of the fusion proteins allows an active luciferase to be formed (18). Here we have used this system to measure dimerization of ACE2 and screened a library of approved drugs (FMC Library (19)) using an unsupervised approach to identify drug candidates for repurposing. Our experiments demonstrated that fenofibric acid (Figure S1), the active metabolite of the oral hyperlipidaemic drug fenofibrate, apparently induced ACE2 dimerization and destabilized the spike RBD inhibiting binding of spike-RBD to ACE2. Importantly, and as hypothesised, fenofibrate-induced changes in RBD-ACE2 interactions correlated with significantly lower infection levels (<60%) in cell culture models using live SARS-CoV-2. Our data combined with unpublished data from other groups and the existing clinical knowledge of fenofibrate identify it as a strong candidate for treating SARS-CoV-2 infections.

## Methods

### Materials

The plasmid pcDNA3 encoding ACE2 was obtained from GenScript (OHu20260); the plasmid encoding prolactin (PRL) was obtained from Sino Biological (HG10275-CY). Optimem and Lipofectamine 2000 were obtained from Thermo Fisher Scientific. NanoBIT and HiBIT detection reagents, Flexicloning transfer systems (C8820 and C9320) and NanoBIT starter kit (N2015) were obtained from Promega. Anti-His antibody was from Thermo Fisher Scientific (37-2900) and Anti-FLAG from Cell Signalling Technology (#2368). The plasmid pcDNA3 encoding ACE2-Flag was obtained from GenScript (OHu20260) and pcDNA3 encoding ACE2-SBP-6xHis was obtained from Thermo Fisher Scientific.

### Molecular biology

Full length ACE2 was amplified by PCR using primers (forward GACCGCGATCGCCATGTCAAGCTCTTCCTGGCTCCTTCT; reverse GATGGTTTAAACAAAGGAGGTCTGAACATCATCAGTG) to introduce a 5’ Sgf1 restriction site immediately prior to the start codon and a Pme1 restriction site directly after the codon encoding the last Phe residue. The PCR product was digested with flexiblend (Sgf1 and Pme1), gel purified and ligated into pF4ACMV before verifying by sequencing. The insert was subsequently transferred into either pFC34K (encoding LgBIT) or pFC36K (encoding SmBIT) using the C-terminal flexicloning system to generate C-terminal fusion proteins.

### NanoBIT assay

HEK-293 cells were grown in DMEM supplemented with 10% (v/v) fetal calf serum and penicillin-streptomycin (50 U/ml). For each well of a 384 plate, 1.25 μl of Optimem containing 10 ng/μL of each of pFC34K ACE2 and pFC36K ACE2 was mixed with an equal volume of Optimem containing 8% lipofectamine-2000. After incubating at room temperature for 30 minutes, the transfection mix was mixed with 10 volumes of well dispersed HEK-293 cells (300,000 cells/mL) in 10% FCS/DMEM without antibiotics and, 25 μL plated per well of white 384 well plates. The two outer rows of the plate were filled with 25 μL media as a humidity barrier. After 48 hours, 2.8 μL drug at 10 x the final concentration were added per well and incubated for 1 hour. Detection reagent was prepared by mixing per well 6.33 μL of detection reagent buffer, 0.33 μL of substrate and 8.34 μL of Optimem containing 10 mM Hepes prewarmed to 37°C. 15 μL detection reagent was added per well, gently mixed and luminescence read every 10 minutes over 30 minutes.

To test whether the drugs inhibit nanoluc directly, HiBIT-RBD was prepared as described previously (12) and the drug added to the desired final concentration, mixed with an equal volume of HiBIT detection reagent and luminescence measured. The results were compared to the luminescence measured using HIBIT-RBD containing DMSO.

To measure whether the drugs inhibited the binding of HiBIT-RBD to ACE2, drugs were tested in the binding assay as previously described on ice (12), Alternatively, binding was measured after 20 min at 37°C.

### Precipitation of ACE2 complexes

HEK-293 cells were transfected by mixing (for each well of a 6 well plate) 0.5 μg of each pcDNA3 ACE2-Flag and pcNDA3 ACE2-SBP-6xHis in 50 μL Optimem. pCMV3 Prolactin (PRL) was used as a negative control in the absence of plasmids encoding ACE2. 50 μL of 8% Lipofectamine-2000 in Optimem was added to plasmid DNA and after 30 min incubation, 1 mL of HEK-293 cells (300,000 per ml) added and the suspension plated per well in 6 well plates. After 12 hours incubation, the cell culture supernatant was gently removed and replaced with fresh DMEM containing 10% FCS. After a further 6 hours, the medium was again removed and the cells lysed in RIPA 250 μL as previously described (20). Lysates were cleared by centrifugation (20,000*g*, 10 min, 4 °C), 30 μL saved for analysis, whilst 200 μL was mixed with 20 μL of streptavidin beads for 2 hours at 4°C. The beads were washed twice with RIPA and once with Tris-buffered saline before being separated on a 4-12% SDS-PAGE gel, transferred to PVDF and proteins detected with anti-FLAG (1/1000) or anti-His (0.08 μg/ml) antibodies.

### Expression of the Spike S1-Receptor Binding Domain for ELISA

Secreted Spike S1 Receptor Binding Domain (RBD) was produced stably using CHOZN GS-/-cells in suspension employing a plasmid encoding residues 319-591 of 2019-nCoV S (upstream of a C-terminal HRV3C protease cleavage site, mFc tag and 8xHis Tag; gifted by Jason S. McLellan, University of Texas, Austin), as described by Tree et al (2020). Coding region of RBD-Fc was subcloned into a modified pCGS3 (Merck/formally known as Sigma-Aldrich) for glutamine selection in CHOZN GS-/- cells. Briefly, RBD-Fc stable clone was obtained by electroporation with 2×10^6^ cells and 5 μg endotoxin-free plasmids using Amaxa kit V and program U24 with Amaxa Nucleofector 2B (Lonza, Switzerland). Electroporated cells were subsequently plated in 96-wells at 500 cells/well in Plating Medium containing 80% EX CELL® CHO Cloning Medium (Cat.no C6366) and EX-CELL CHO CD Fusion serum-free media without glutamine. High expressing clones were scaled-up in serum-free media without L-glutamine in 50 mL TPP TubeSpin® shaking Bioreactors (180 rpm, 37°C and 5% CO2) for RBD-Fc production. A HiTrap Protein G, HP column (GE Healthcare, US), equilibrated in 1x PBS prior to use, was employed to purify the Spike S1 RBD, eluting with glycine (100 mM, pH 2.7). Purity was confirmed using SDS-PAGE with Coomassie stain and quantified using the bicinchoninic acid assay (Thermo Scientific).

### ELISA assay measuring RBD-ACE2 binding

An RBD-ACE2 inhibition ELISA was performed as described by Tree *et al* (2020). Streptavidin (3 μg/mL; Fisher) was precoated onto the surface of 96 well plates (high binding; Greiner) in Na_2_CO_3_ buffer (50 mM; pH 9.6; 1 hour; 37°C). Plates were washed 3x (300 μL PBS containing 0.2% w/v Brij35) prior to blocking for 1 hour at 37°C with 50 μL PBS, 0.2% w/v Brij35, 1% w/v casein. After washing 3x with PBS, plates were coated with 50 μL of 100 ng/mL biotin-ACE2 (Sino Biological) in PBS containing 0.2% w/v Brij35, 1% w/v casein for 1 hour at 37°C. Plates were then washed and incubated at room temperature in 50 μL of 5 μg/mL RBD in PBS containing 0.2% w/v Brij35, 1% w/v casein for 30 minutes in the presence or absence of test drugs. Plates were incubated (1 hour; 37°C) to allow binding before 3 washes. Bound RBD was detected by incubation (1 hour; 37°C) with rabbit anti-SARS-CoV-2-Spike-RBD (Stratech) (1:2000 v/v in PBS containing 0.2% w/v Brij35, 1% w/v casein. Following 3 further washes, plates were incubated (30 mins.; at 37°C) with horseradish peroxidase-conjugated donkey anti-rabbit IgG (1:2500 v/v, in PBS containing w/v Brij35, 1% w/v casein. Plates were washed five times before the addition of 3,3’,5,5’-tetramethylbenzidine substrate, prepared as per manufacturer’s instructions (Sigma-Aldrich). Colour development was halted after 10 mins by the addition of H_s_SO_4_ (2 M) and quantified at λ_abs_ = 450 nm using a Tecan Infinite M200 Pro multi-well plate reader (Tecan Group). Specific binding was determined by subtracting the absorbance measured in samples lacking ACE2.

### Differential scanning fluorimetry

Differential scanning fluorimetry (DSF) was conducted with 1 μg RBD in 40 μL PBS (pH 7.6) with 1.25x SYPRO™ Orange (Invitrogen) and either, H_2_O, sodium acetate or fibrates in 96-well qPCR plates (AB Biosystems). An AB Biosystems, StepOne plus, qPCR machine with a TAMRA filter was employed to perform melt curve experiments, increasing the temperature by +0.5°C every 30 seconds, from 25 - 90°C. First-order differential plots were calculated after smoothing (Savitzky-Golay, 9 neighbours, 2^nd^-order polynomial) using Prism 8 (GraphPad). The peak maxima of the first-order differential plots were determined with MatLab software (R20018a, MathWorks) and used to calculate the change in T_m_ in the presence of fibrates. Control wells without RBD, but containing sodium acetate or fibrates, were tested to confirm that altered T_m_ values were a result of protein-ligand interactions and not a result of an interaction between the drug and the dye.

### SARS-CoV-2 infection experiments (hCOV-19/England/2/202 strain)

Vero cells (ATCC® CCL-81) were washed with PBS, dislodged with 0.25% Trypsin-EDTA (Sigma life sciences) and seeded into 96-well imaging plates (Greiner) at a density of 8×10^3^/well in culture media (DMEM containing 10% FBS, 1% Penicillin and Streptomycin, 1% L-Glutamine and 1% non-essential amino acids). The next day, cells were infected with SARS-CoV-2 strain hCOV-19/England/2/2020, isolated by Public Health England (PHE) from the first patient cluster in the United Kingdom on 29 January 2020. Virus stock 10^6^ IU/ml (kind gift from Christine Bruce, Public Health England) was diluted 1/150 in culture media allowing 25 μl per well. Virus was then diluted further with 25μl per well media containing treatments of interested prepared at 2X concentration to give 1x drug and a final virus dilution of 1/300. Cells were then infected with virus (167 IU/well) and cultured for 24 or 48 hr. After the infection period, supernatants were harvested and frozen prior to analysis by qRT-PCR, and cells were fixed in ice-cold methanol. Cells were then blocked in PBS containing 10% FBS and stained with rabbit anti-SARS-CoV-2 spike protein, subunit 1 (The Native Antigen Company), followed by Alexa Fluor 555-conjugated goat anti-rabbit IgG secondary antibody (Invitrogen, Thermo Fisher Scientific). Cell nuclei were stained with Hoechst 33342 (Thermo Fisher Scientific). After washing with PBS, cells were imaged and analysed using a Thermo Scientific CelIInsight CX5 High-Content Screening (HCS) platform. Infected cells were scored by perinuclear fluorescence above a set threshold determined by positive (untreated) and negative (uninfected) controls. A minimum of 9 fields and 5000 nuclei per well in triplicate or quadruplicate wells per treatment were scored in each experiment. All experiments were performed 2-4 times.

### SARS-CoV-2 plaque formation assay (Italy/UniSR1/2020 strain)

Vero cells were plated at 2.5 x 10^5^ cell/well in 24-well plates in Essential-modified Eagle Medium (EMEM, Lonza) supplemented with 10% fetal calf serum (FCS, EuroClone) (complete medium). Twenty-four hours later, cells were incubated with compounds in 250 μl of complete medium 1 hour prior to infection and then incubated with virus suspension (pre-treatment) containing 50 plaque forming units (PFU) of Italy/UniSR1/2020 strain (GISAID accession ID: EPI_ISL_413489). After incubation for 1 hour at 37°C, supernatants were discarded, 500 μl of 1% methylcellulose (Sigma Chemical Corp) overlay dissolved in complete medium was added to each well. Alternatively, Vero cells were incubated with compounds together with a virus suspension containing 50 PFU (co-treatment) in a total volume of 300 μl complete medium for 1 hour. Supernatants were discarded and the methylcellulose overlay was added as described above. After 3 days, cells were fixed using 6% formaldehyde/PBS solution for 10 minutes and stained with 1% crystal violet (Sigma Chemical Corp) in 70% methanol for 1 hour. The plaques were counted under a stereoscopic microscope (SMZ-1500, Nikon).

### Quantitative Real time PCR for SARS-CoV-2

Cell culture supernatant from infection experiments was heat-inactivated at 56°C for 60 mins following PHE protocols in the NHS Turnkey Labs based in the University of Birmingham Medical School. Viral RNA was reverse transcribed and quantified in culture supernatant using the 1-step SARS-CoV-2 Viasure Real Time PCR Detection Kit (Prolab Diagnostics/CerTest Biotec) according to manufacturer’s instructions. Briefly, 15μl of rehydrated Reaction-Mix was combined with 5μl of either heat-inactivated cell culture supernatant, positive virus RNA control or negative control before cycling in an Agilent AriaMX Real-Time thermal cycler using the following cycle conditions: reverse transcription at 45°C for 15 mins, initial denaturation at 95°C for 2 mins followed by 45 cycles of 95°C for 10 sec, 60°C for 50 sec. Fluorimetric data was collected during the extension step for FAM (ORF1ab gene), ROX (N gene) and Hex (internal control) and Cycle thresholds (Ct) calculated for each gene. Relative expression was calculated by subtracting the Virus Control Ct values from drug treatment samples and transforming the data using 2-Δ^Ct^.

### Statistical analysis

All pairwise comparisons were performed using paired T-Tests or Mann-Whitney U tests where normal distribution was not assumed. Multiple comparisons were done using ANOVA.

## Results

### Validation of ACE2 dimerization assay

To develop an assay to measure dimerization of ACE2, two separate plasmids were created encoding ACE2 fused in frame at its C terminus to one of the nanoBIT reporters, SmBIT or LgBIT (Figure 1A). When these constructs were expressed in HEK293 cells, luminescence was observed that was approximately 20% of that generated by expression of LgBIT and SmBIT fused to the protein kinase A regulatory (PRKAR2) and catalytic (PRKACA) subunits, respectively (positive control). Co-transfection of plasmids encoding ACE2 fused to either LgBIT or SmBIT and PRKAR2 or PRKACA subunits fused to the complementary nanobit reporter did not generate luminescence, suggesting that the assay measured ACE2 dimerization (Figure 1B). Luminescence was also not observed when cells were transfected with nanoBIT-tagged ATG5 and PRKAR2, two proteins known not to interact (Figure 1B). To confirm the assay measured ACE2 dimerization, cells were transfected with a plasmid encoding untagged ACE2 as well as ACE2 tagged with LgBIT or SmBIT. The untagged ACE2 was expressed under the control of a CMV promoter, which provides substantially higher-level expression than the HSV TK promoter which controls the expression of the NanoBIT-tagged ACE2. If the assay measures dimerization, expression of the untagged ACE2 would be expected to suppress the luminescence by competing with the tagged ACE2 in dimers. To ensure the effect observed did not result from competition for transcription factors, rather than as a result of the untagged ACE2 competing with NanoBIT tagged ACE2, an unrelated gene (prolactin-PRL) was also expressed under the control of the CMV promoter. High level expression of untagged ACE2 suppressed the luminescence signal generated by ACE2 tagged with the NanoBIT reporters but it did not suppress the luminescence measured with the NanoBIT-tagged protein kinase A subunits (Figure 1C).

**Figure 1.**
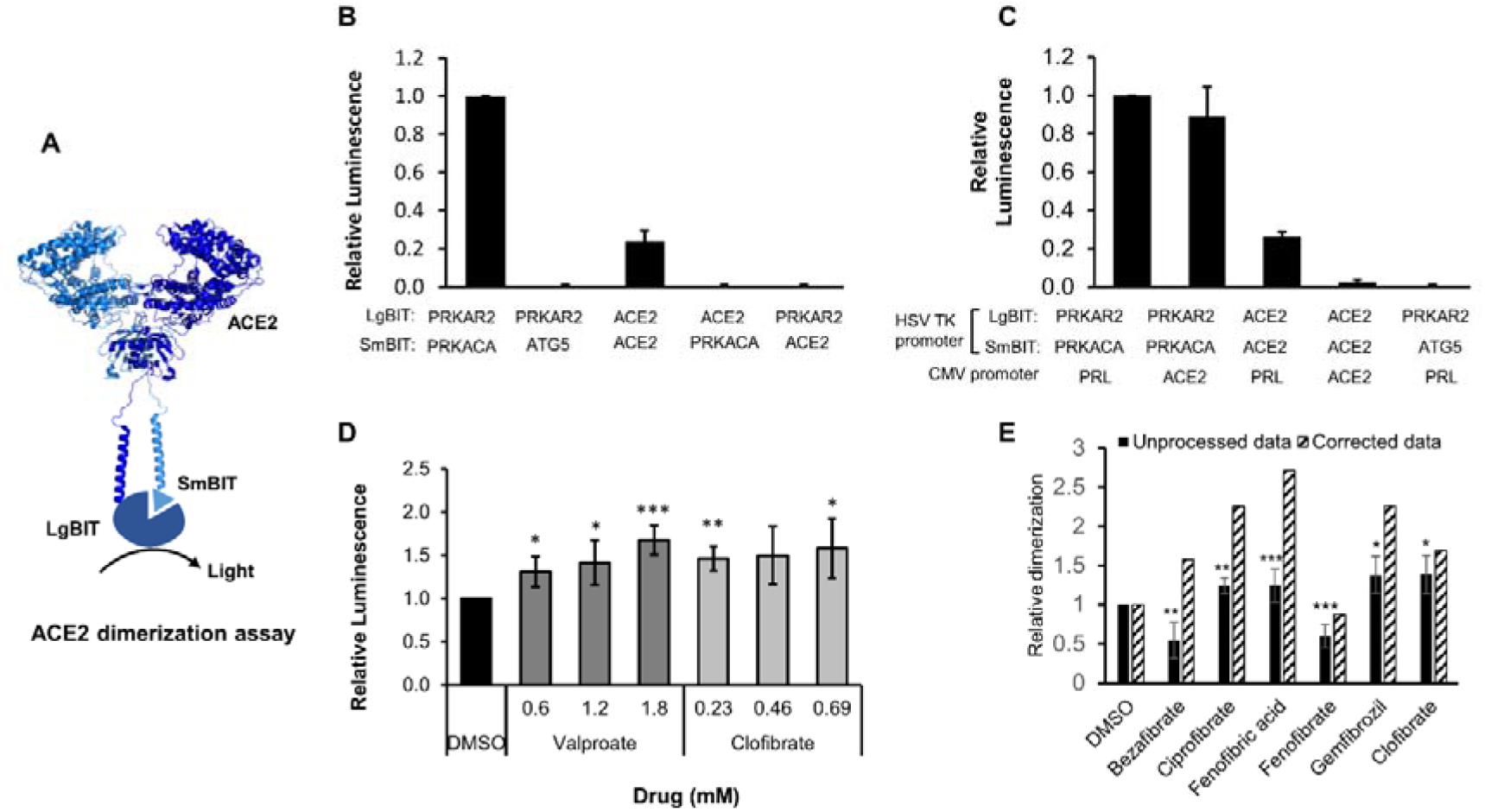
ACE2 dimerization assay. **A.** Schematic showing ACE2 tagged with LgBIT and SmBIT. **B.** HEK-293 cells were transfected with combinations of plasmids encoding LgBIT or SmBIT fused to either protein kinase A regulatory subunit (PRKAR2) or catalytic subunit (PRKACA), ATG5 or ACE2. The results (mean ± S.D., n = 5) were normalized to the luminescence measured in cells transfected with protein kinase A reporters (positive control). **C.** HEK-293 cells were transfected with plasmids encoding ACE2 nanoBIT reporters under the control of the HSV TK promoter and ACE2 or prolactin (PRL) under the control of the CMV promoter. The results (mean ± S.D., n = 4) were normalized to the luminescence measured in cells transfected with protein kinase A reporters and prolactin. **D.** HEK-293 cells were transfected with NanoBIT-tagged ACE2 reporters and incubated with sodium valproate or clofibrate at a concentration equal to 1x, 2x or 3x the reported C_max_ of the drug. After 1 hour, luminescence was measured and normalized (mean ± S.D., n =4) to that measured in cells treated with DMSO. **E.** A series of other fibrates were similarly evaluated in the assay. The luminescence measured (mean ± S.D., n = 5-11, solid bars) was significantly different to that measured in cells treated with solvent where shown. When these fibrates were incubated with purified LgBIT and HiBIT-RBD to create a constitutively active nanoluc, each of these fibrates were found to inhibit nanoluciferase (bezafibrate 35 ± 7 %, ciprofibrate 55 ± 6 %, fenofibric acid 46 ± 3 %, fenofibrate 69 ± 5 %, gemfibrozil 61 ± 2 % of the activity measured in the presence of DMSO). To correct for this, the luminescence measurements from cells treated with fibrates in cells were divided by these latter values to estimate the effect of the drugs on dimerization (hatched bars). Significant difference from control is shown as *, *P*< 0.05; **, *P* < 0.01; ***, *P*< 0.005.

### Identification of ACE2 dimerization modulators

The assay was used to screen a custom in-house library of approximately 100 approved drugs at a final concentration equal to their C_max_ in patients (FMC1 Library (19)). Sodium valproate and clofibrate both increased the dimerization signal by approximately 33% and 56%, respectively. To confirm this, fresh compounds were purchased and retested at a concentration equal to their C_max_ in patients and multiples of this. Both compounds significantly increased the measured luminescence, confirming the results of the screen (Figure 1D). Although clofibrate has previously been approved, it has subsequently been withdrawn due to unacceptable toxicity (21). However, several other fibrates are still in clinical use. Apart from fenofibrate, these all bear a carboxylic acid whereas fenofibrate is an isopropyl ester pro-drug of fenofibric acid (Figure S1). Noting that sodium valproate is also a lipophilic carboxylic acid, fenofibric acid was tested in the dimerization assay. All of the fibrates (tested at 230 μM, the C_ss_ of clofibrate (22)) modestly, but significantly, increased luminescence (Figure 1E). However, they also substantially decreased the luminescence generated by mixing LgBIT with HiBIT-tagged RBD (which binds LgBIT with high affinity and independently of other interacting molecules). This suggested that the drugs inhibited nanoluc directly and the measured luminescence underestimated dimerization. When the luminescence measured in the assay was corrected to take into account inhibition of nanoluciferase (Fig 1E, corrected data), fenofibric acid emerged as the most effective, apparently increasing dimerization by approximately two-fold. In contrast to this, fenofibrate did not increase the dimerization. The increase in luminescence was also time-dependant, reaching a maximum after 30 minutes exposure to the drug (Figure S2).

To confirm these results, HEK-293 cells were transfected with plasmids encoding ACE2 tagged with streptavidin binding protein and a His-tag or ACE2 with a FLAG tag. Cells were exposed to drug, lysed and ACE2 complexes purified using streptavidin beads. Following immunoblotting, ACE2-Flag was only detected in lysates from cells transfected with both plasmids and not from cells transfected with one plasmid alone, confirming the assay measured the interaction of ACE2. However, when cells were exposed to the fibrates, the amount of ACE2-FLAG detected on the beads was not substantially altered (Figure S3).

### Effect of fibrates on S protein RBD

To evaluate whether fibrates affect the viral spike protein RBD, the thermal stability of RBD in the presence and absence of fibrates was investigated using differential scanning fluorimetry (DSF). Changes in the T_m_ of a protein in the presence of a ligand is indicative of binding and has previously been utilised to probe for protein-ligand interactions (23). All of the fibrates altered the T_m_ of RBD (46.4°C) although the greatest destabilization was observed with bezafibrate and ciprofibrate (both ΔT_m_ =-1.9°C, (Figure S4). A smaller effect was observed with fenofibric acid (ΔT_m_ =-1.4°C) but this was detectable at concentrations as low as 30 μM (Figure 2A, B). Although fenofibrate also destabilized RBD, this was only observed at higher drug concentrations (≥ 270 μM, Figure S4). Acetate, a carboxylic acid lacking the lipophilic moieties found in the fibrates, had no significant effect on RBD T_m_ (Figure S4) indicating that the lipophilic moieties are required.

**Figure 2.**
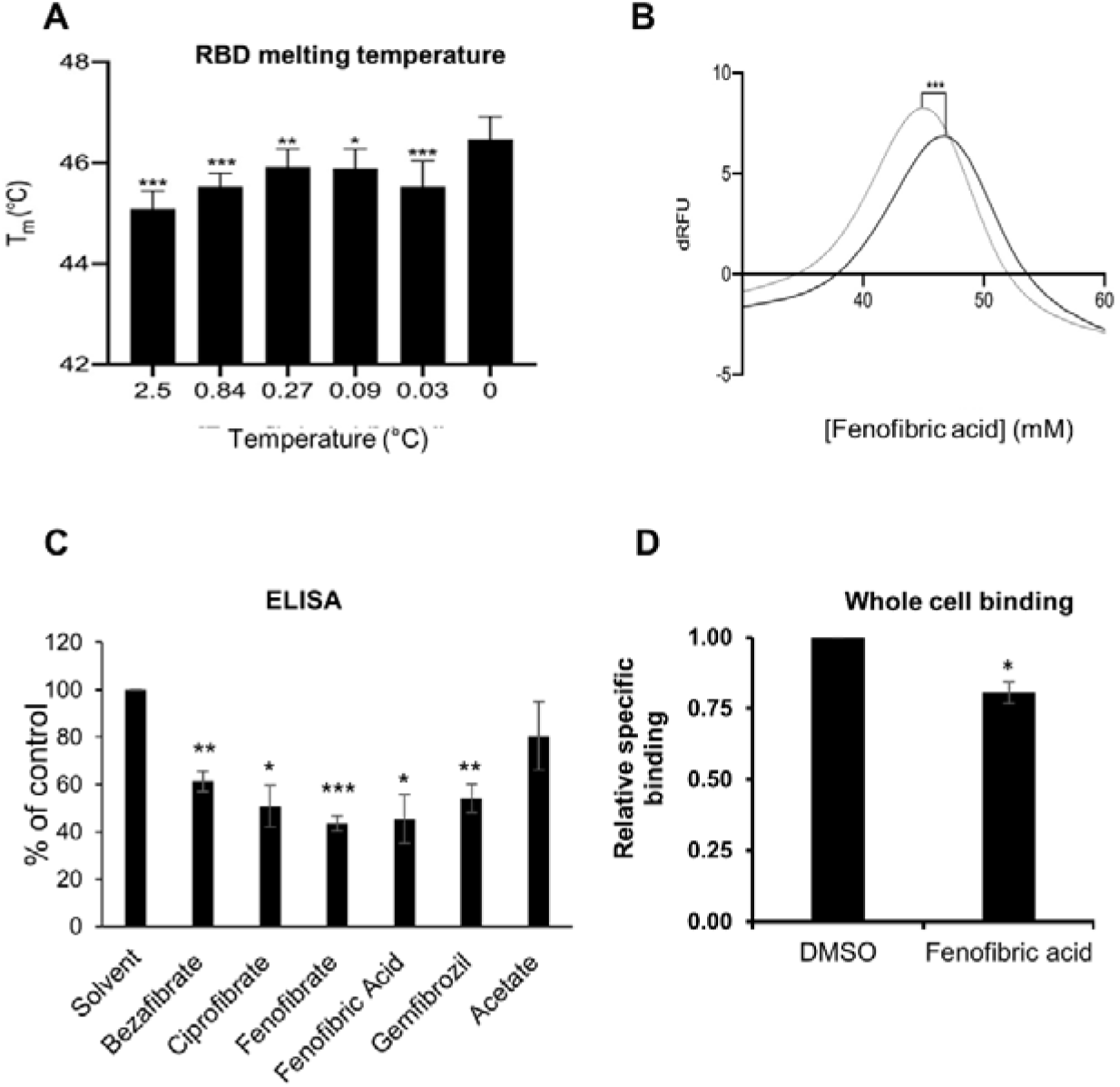
Effect of fenofibrates on RBD and RBD binding to ACE2. **A.** Differential scanning fluorimetry. The T_m_ of 1 μg RBD alone or with increasing concentrations of fenofibric acid. The results (mean ± S.D., n=3) were significantly different from RBD where shown (***, *P* < 0.001; **, *P* < 0.01; *, *P* < 0.05; paired t-test). **B.** First differential of the thermal stability of 1 μg RBD alone (solid line) or with 2.5 mM fenofibric acid (dotted line). A direct interaction of fenofibric acid with SYPRO™ Orange dye (in the absence of RBD) was not observed. **C**. ELISA assay to measure inhibition of RBD binding to ACE2 by Fibrates. Biotinylated ACE2 was captured onto a high binding microplate coated with streptavidin prior to the addition of RBD pre-incubated with or without 230 μM bezafibrate, ciprofibrate, fenofibrate, fenofibric acid, gemfibrozil or acetate control. Data (mean ± S.D., n=3) represented as % no inhibitor control and are significantly different to this where shown (*, *P* < 0.05; **, P<0.01; ***, P<0.005). **D**. A Whole cell binding assay to measure inhibition of RBD binding to ACE2. COS cells were transfected with ACE2 and incubated on ice with HiBIT-tagged SARS-CoV-2 RBD and the indicated fibrate (230 μM). After washing, the bound RBD was measured by addition of LgBIT and nanoluc substrate. The results mean ± S.D., n = 4) were normalized to the binding measured in cells exposed to DMSO and are significantly different where shown (*, *P* < 0.005).

### Fenofibric acid inhibits ACE2-RBD binding

An ELISA assay consisting of immobilised, recombinant ACE2 was employed to determine the inhibitory effect of fibrates on RBD-ACE2 binding. All fibrates screened demonstrated significant inhibition of binding at a concentration of 230 μM, the C_max_ of clofibrate (Figure 2C). The binding of RBD to ACE2 expressed in COS cells was measured as previously described (12). When these assays were conducted on ice to minimize endocytosis, no inhibition of RBD binding was observed with any of the fibrates (Figure S5). However, when the assay was adapted for use at 37°C by using shorter incubation times, fenofibric acid was found to modestly, but significantly, inhibit RBD binding to ACE2 (Figure 2D). This was not due to toxicity as 99 ± 1% (n=4) of the cells excluded trypan blue after a similar exposure to drug. Furthermore, in a preliminary experiment, fenofibric acid inhibited binding to fixed Vero cells (Figure S5). Combined, these data indicate that fenofibrate/fenofibric acid interfere with spike RBD binding to ACE2.

### Fenofibrate inhibits infection of Vero cells by the hCOV-19/England/2/2020 virus isolate

To evaluate the potential therapeutic effect of fenofibrate/fenofibric acid on SARS-CoV-2 virus, infection experiments were performed independently in two separate laboratories. Using the hCOV-19/England/2/2020 virus strain, Vero cells were co-incubated with virus and fibrates before fixing and staining for spike protein and counterstaining nuclei with Hoescht. Using live virus allows measurement of both primary infection after 24 hours by the viral inoculum and subsequent reinfection by virus released by Vero cells in the wells (after 48 hours). By 48 hours, 59% of Vero cells stained positive for spike protein in virus control wells with minimal loss of cell numbers (Figure 3 A & B, Figure S6). Consistent with the binding assays, and of the fibrates studied (all screened at 230 μM), only fenofibrate reduced virus infection by ~65% to 18% compared to virus control (Figure 3B, Figure S5). This was not attributable to loss of Vero cell viability as no decrease in cell number by fenofibrate was seen as measured by number of nuclei (Figure 3B, Figure S5) and by Cell Titre Blue assay (Figure S8). No difference was observed when cells were pretreated or co-treated with drug and virus (data not shown). Parallel experiments were performed with a panel of statins (simvastatin, pitavastatin, rosuvastatin and pravastatin, Figure S1), drugs which have largely replaced fibrates as front-line therapy for reducing cholesterol levels and treating lipid disorders. When screened at 100 nM, a significant decrease in infection rates was observed with simvastatin and pitavastatin but not with pravastatin or rosuvastatin (Figure 3 C & D). However, this decrease was associated with significant loss of Vero cell viability as measured by decrease in number of nuclei (Figure 3 C & D; Figure S6-S7) and cell titre blue assay. Titration experiments were performed with simvastatin and pitavastatin on Vero cells and viability assessed in the absence of virus. A concentration of 10 nM did not affect Vero cell viability after 48 hours and no reduction in infection was observed (Figure S7) indicating that this panel of statins do not modulate SARS-CoV-2 infection, at least not *in vitro*.

**Figure 3.**
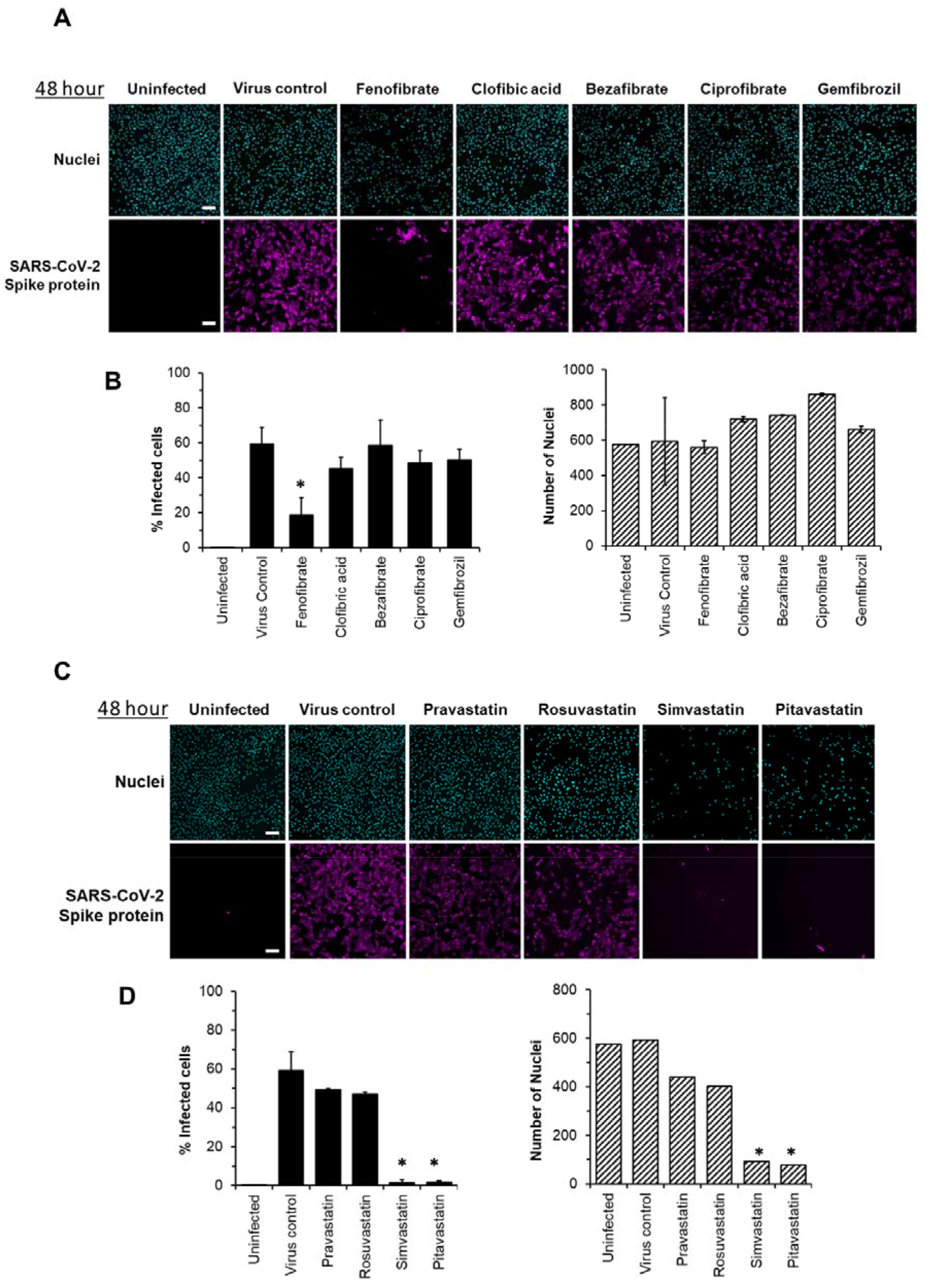
Fenofibrate reduces SARS-CoV-2 infection rates in vitro. Vero cells were plated into 96 well plates (8 x 10^3^ cells/well) for 24 hours before infecting with 167 IU of hCOV-19/England/2/2020 virus isolate in the absence or presence of drugs. Infection rates were assessed at 48 hours by staining Vero cells for viral Spike protein and counterstaining nuclei with Hoescht. Cells were imaged and analysed using a Thermo Scientific Celllnsight CX5 High-Content Screening (HCS) platform. Representative images and mean data are shown for Vero cells incubated with either no virus, SARS-CoV-2 virus control, or virus and fibrates (230μM, (**A and B**) or statins (100nM, **C and D**). The black bars are % infected cells and the hatched grey bars are average number of nuclei score per field of view (mean ± S.D., n=2-3; one-way ANOVA. *, *P* < 0.05 compared to virus control).

Subsequent experiments assessed the effect of fenofibrate and fenofibric acid on infection by SARS-CoV-2. Within 24 hours, fenofibrate had significantly reduced infection levels by ~60% indicating that fenofibrate is able to inhibit primary infection (Figure 4 A & B). A reduction was also observed with fenofibric acid, albeit less than fenofibrate, however the results were more variable in the experiments performed and did not reach significance (Figure 4A-B). This pattern was recapitulated at 48 hours (Figure 4C-D) indicating that suppression of infection by fenofibrate is sustained. These data indicate that fenofibrate, and to a lesser extent fenofibric acid, are able to reduce primary infection and also secondary infection rates.

**Figure 4.**
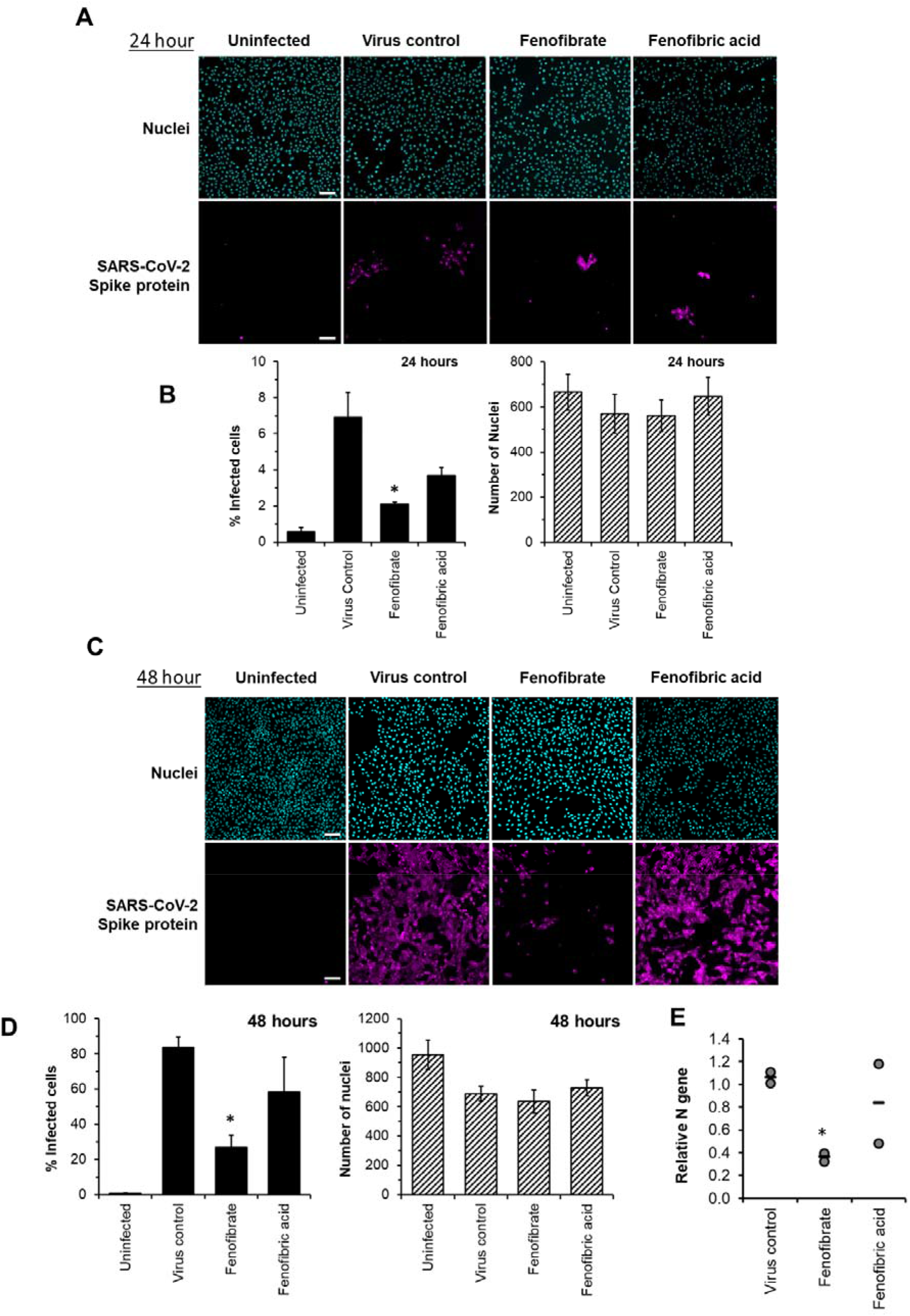
Fenofibrate, and to a lesser extent fenofibric acid, reduce SARS-CoV-2 infection at both 24 and 48 hours. Vero cells were plated into 96 well plates (8×10^3^ cells/well) for 24 hours before infecting with 167IU of hCOV-19/England/2/2020 virus isolate in the absence or presence of 230μM fenofibrate or fenofibric acid. Infection rates were assessed at 24 and 48 hours by staining Vero cells for viral Spike protein and counterstaining nuclei with Hoescht. Cells were imaged and analysed using a Thermo Scientific CellInsight CX5 High-Content Screening (HCS) platform. Representative images and mean data are shown for Vero cells incubated for 24 hours (**A** and **B**) and 48 hours (**C** and **D**). The black bars are % infected cells and the hatched grey bars are average number of nuclei score per field of view (mean ±S.D. n = 2-3 one-way ANOVA, *, *P* < 0.05 compared to virus control). **E.** Supernatant was collected from wells after 48 hours of incubation. Virus was heat-inactivated and viral N gene RNA levels measured directly in supernatant using a commercial one-step RT-qPCR reaction. N RNA levels were calculated relative to supernatant from virus control (n=2 experiments).

To determine virus levels in cell culture supernatant, virus RNA levels were measured by multiplex qRT-PCR for viral ORF1ab and N genes on heat-inactivated culture supernatant from 48 hour experiments. Whilst ORF1ab RNA levels were detectable in virus control supernatant, no signal was detected in supernatant from drug-treated cells implying, but not proving, a reduction in virus RNA (data not shown). However, a signal for the viral N-gene was detectable by qRT-RCR in all samples. Consistent with the reductions seen in infection levels, fenofibrate significantly reduced viral N-gene RNA levels whereas the results with fenofibric acid were more variable (Figure 4E). Furthermore, the effect of fenofibrate on infection rates and viral RNA levels in culture supernatant was dose-dependent as determined by doubling dilution experiments (1x: 230 μM; Figure 5 A & B). Fenofibrate works as an anti-hyperlipidaemia agent by acting as a PPARα agonist. Treatment with the PPAR-alpha antagonist GW6471 did not significantly alter the anti-viral actions of fenofibrate (Figure 5 C &D) suggesting that the antiviral actions of fenofibrate in this system are independent of PPARα.

**Figure 5.**
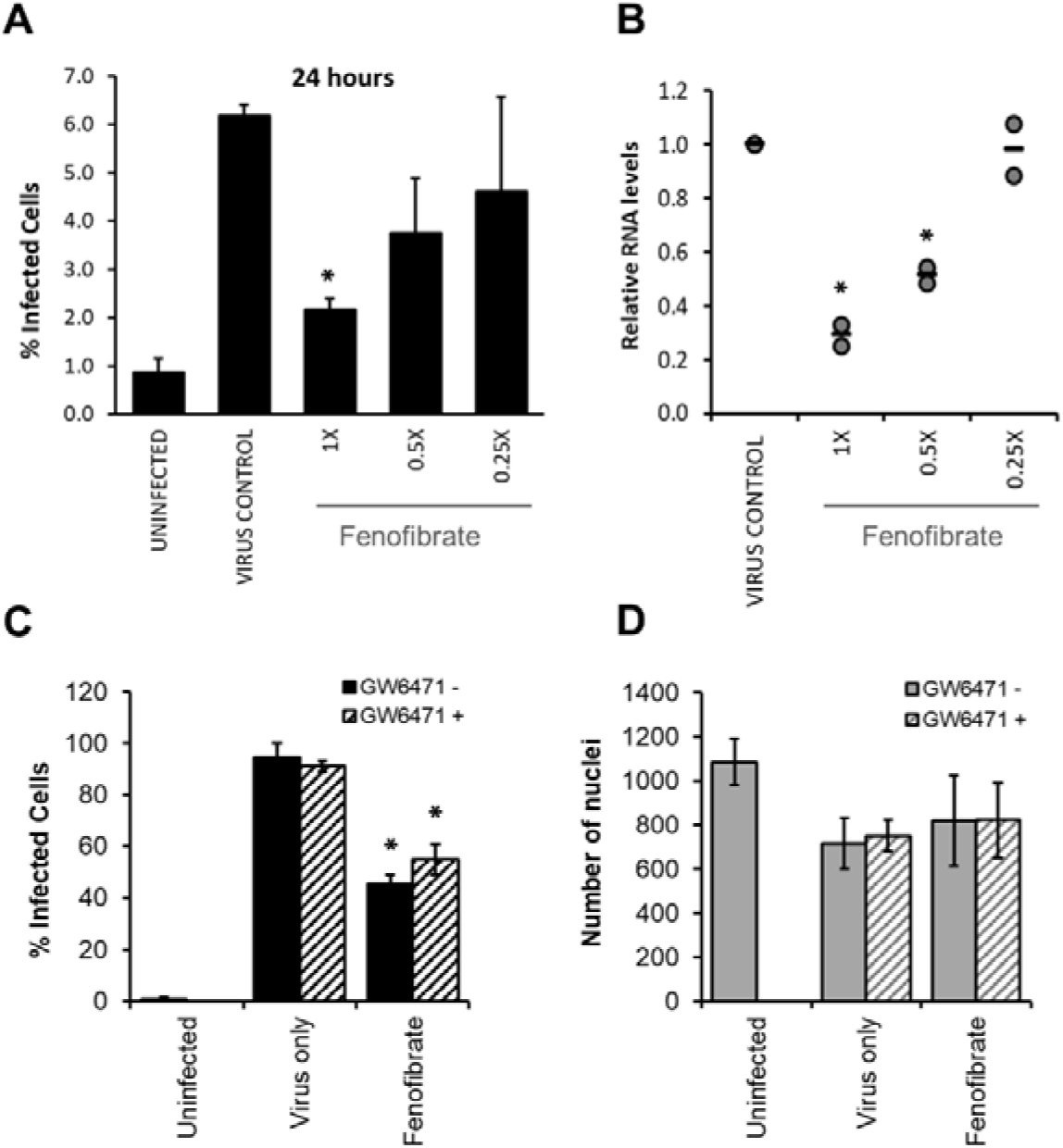
Fenofibrate reduces SARS-CoV-2 infection level in vitro in a dose dependent manner. Vero cells were plated into 96 well plates (8 x10^3^ cells/well) for 24 hours before infecting with 167 IU of hCOV-19/England/2/2020 virus isolate in the absence or presence of 1x (230μM), 0.5x or 0.25x fenofibrate. Infection was assessed at 24 hours by staining Vero cells for viral Spike protein and counterstaining nuclei with Hoescht. Cells were imaged and analysed using a Thermo Scientific CelIInsight CX5 High-Content Screening (HCS) platform. (**A**) Mean infection rates observed at 24 hours (n=2-3). **B.** Supernatant was collected from wells after 48 hours of incubation. Virus was heat-inactivated and viral N gene RNA levels measured directly in supernatant using a commercial one-step RT-qPCR reaction. N RNA levels were calculated relative to supernatant from virus control (n=2). To determine the role of PPARα, 48 hour infection experiments were performed in the absence or presence of the PPAR-alpha antagonist GW6471 (1 μM). Mean data from 2-3 experiments are shown in (**C** and **D**). **C** shows % infected cells and **D** the average number of nuclei score per field of view. Statistical significance was calculated by one-way ANOVA. *, *P* <0.05 compared to virus control.

### Fenofibrate inhibits infection of Vero cells by the Italy/UniSR1/2020 virus isolate

To confirm the infection results observed with hCOV-19/England/2/2020 isolate in experiments performed at the University of Birmingham, the effect of fenofibrate and fenofibric acid was assessed on plaque formation in Vero cells infected with the Italy/UniSR1/2020 SARS-CoV-2 isolate independently at San Raffaele Scientific Institute in Milan. Vero cells were pretreated for 1 hour with fenofibrate or fenofibric acid or were exposed to the drug and the virus at the same time (co-treatment). Fenofibric acid inhibited plaque formation at concentrations clinically achievable in patients. The reduction of plaque formation by fenofibric acid reached 62% at 50μM drug in the co-treatment condition (Figure 6). Fenofibrate also reduced the number of plaques formed, but notably less potently. As observed for the hCOV-19/England/2/2020 strain, no difference was observed between pre-treatment and co-treatment experiments.

**Figure 6.**
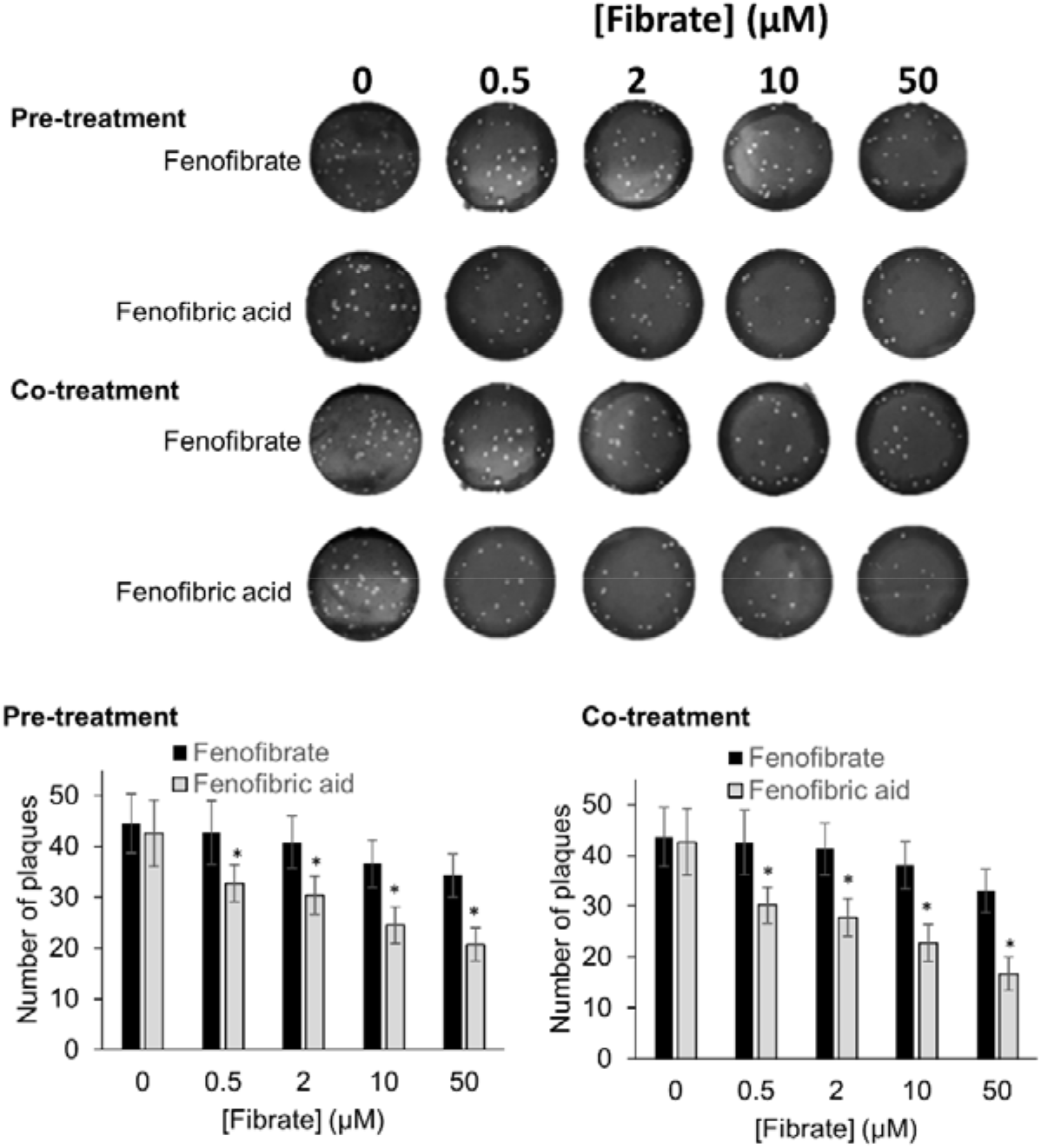
Fibrate inhibition of SARS-COV-2 infection of Vero cells. Antiviral effect of fibrates added 1 hour before infection or in co-treatment with infection in Vero cells with 50 PFU of SARS-CoV-2. N.D, not determined due to solubility issues. The results are expressed as number of PFU/well and represent the mean ± SD of two experiments each with 3 separate plates containing duplicate samples. The number of plaques was significantly different (2 way Anova) in cells treated with fenofibric compared to fenofibrate where shown DMSO (*, *P* < 0.001). Compared to cells treated with drug solvent, the number of plaques was significantly different in cells treated with fenofibric acid (*P* < 0.001, all concentrations tested) and in cells treated with fenofibrate (*P* < 0.01, 10 μM fenofibrate; *P* < 0.001, 50 μM fenofibrate).

Thus, using two different virus isolates, we demonstrate that fenofibrate, or its active metabolite fenofibric acid, are able to significantly reduce SARS-CoV-2 infection in cell culture models.

## Discussion

The development of new more infectious SARS-CoV-2 variants has resulted in a rapid expansion in infection rates and deaths in several countries around the world, especially the UK,US and Europe. Whilst vaccine programmes will hopefully reduce infection rates and virus spread in the longer term, there is still an urgent need to expand our arsenal of drugs to treat SARS-CoV-2-positive patients. Using an unsupervised approach, we have identified that the off-patent licensed drug fenofibrate has the potential to treat SARS-CoV-2 infections. The drug was identified through a screen of approved drugs to identify those which alter dimerization of ACE2. Clofibrate was identified as a hit in this screen and testing of other fibrates led to the identification of fenofibrate as being the most likely to be effective as an antiviral agent. Fenofibrate also appears to affect the stability of spike protein RBD and inhibit binding to ACE2. Importantly, these effects on RBD by fenofibrate correlated with decreases in SARS-CoV-2 infection rates *in vitro* using two different virus assays (staining for Spike protein and plaque-formation) in two independent laboratories.

The ACE2 dimerization assays depends on the co-localization of LgBIT and SmBIT brought about by the formation of ACE2 dimers. No signal was observed using protein kinase A subunits that do not interact with ACE2 and over-expression of unlabelled ACE2 suppressed the signal from the nanobit reporters, giving confidence that the assay measures the interaction of ACE2 protomers. Although described here as a dimerization assay, the assay may not discriminate between dimer formation and higher-order oligomers, and drugs showing activity in the dimerization assay could alternatively elicit conformational changes in ACE2 complexes which improve the interaction of the nanobit reporters. All the fibrates tested showed some activity in the dimerization assays, but the most pronounced effects were observed with fenofibric acid. The pro-drug fenofibrate (the isopropyl ester of fenofibric acid) was inactive in this assay, suggesting the free carboxylic acid is necessary.

In addition to effects on ACE2, all the fibrates destabilized the viral spike protein RBD and lowered its “melting” temperature. However, the most potent effects were again seen with fenofibric acid. This may contribute to fenofibrate inhibiting binding of RBD to ACE2 in ELISA and cell binding studies performed at 37°C. When measured in cells at 0°C, the fibrates did not inhibit binding to ACE2; this temperature is likely to prevent melting, providing a potential explanation for the lack of activity at fibrates in binding assays at lower temperatures. Blocking RBD binding to ACE2 was anticipated to reduce infection by SARS-CoV-2.

To provide robust data evaluating the potential of fenofibric acid/fenofibrate to inhibit infection by SARS-CoV-2, the drugs were evaluated independently in two separate laboratories using different assays and two different SARS-CoV-2 isolates (hCOV-19/England/2/2020 and Italy/UniSR1/2020). In both cases, fenofibrate/fenofibric acid were found to significantly reduce infection rates. Fenofibrate/fenofibric acid decreased the number of Vero cells staining positive for viral spike protein at 24 hours indicating inhibition of primary infection. The number of cells infected 48 hours after infection was also significantly reduced, demonstrating the potential for sustained inhibition of infection. This was further confirmed by PCR which showed a reduction in viral mRNA released by the cells into the culture supernatant. Likewise, we saw significant reductions with fenofibric acid/fenofibrate in plaque formation assays which are considered the gold-standard assay for measuring infectivity by SARS-CoV-2. Several assays demonstrate that the reduced viral infection was not due to a cytotoxic effect of the fibrates in the host cells. Considering that fenofibrate is used in the treatment of hypercholesterolaemia and hyperlipidaemia, the effect of several statins on SARS-CoV-2 infection was also assessed. These included both hydrophilic (pravastatin, rosuvastatin) and lipophilic statins (pitavastatin, simvastatin). None of the statins inhibited viral infection, suggesting the anti-viral effect was not mediated by inhibition of cholesterol synthesis. The differences we observed in potency between fenofibrate and fenofibric acid in the two antiviral assays may reflect different strains of the virus or different methodologies. Although we cannot presently fully explain these, it is clear that fenofibrate or its metabolite fenofibric acid demonstrated anti-SARS-CoV-2 activity.

Fenofibric acid was identified as a potential anti-viral agent through its effects on ACE2 dimerization, but it remains to clarified to what extent the effects of fenofibrate/fenofibric acid on dimerization contribute to its anti-viral activity. The mechanism by which increased dimerization could inhibit viral infection was not investigated and several explanations are plausible. It was not possible to measure an effect of fibrates on dimerization of ACE2 in streptavidin precipitation assays. This may reflect the insensitivity of this latter method or that fenofibrate alters the conformation of ACE2 rather than inducing dimerization. Structural studies have shown that ACE2 adopts “open” and “closed” conformations (13) which may be detected by the nanobit reporters. The open and closed conformations may also affect RBD binding to each ACE2 protomer or the number of spike proteins that can bind to an ACE2 dimer, thereby affecting the avidity of the virus for cells. Conformational changes in ACE2 may also affect its susceptibility to proteolysis by TMPRSS2. The suggestion that the anti-viral activity of fenofibrate depends at least in part on effects on ACE2 also offers advantages over drugs which inhibit viral proteins. Mutations in the viral genome are less likely to affect the antiviral activity of drugs which target human rather than viral proteins. Excitingly, fenofibrate also destabilized the RBD and reduced binding of it to ACE2. It is highly likely that this contributes to the reduced infection in cells treated with fenofibrate. This also suggests that fenofibrate has multiple mechanisms of action, making it less likely that resistance to it will quickly emerge and fenofibrate may retain activity against newly emerging strains of SARS-CoV-2. However, our data suggest that the antiviral activity of fenofibrate measured in the infection assays presented here is not mediated by the transcription factor PPARα. The efficacy of fibrates in the treatment of hyperlipidaemia depends on their ability to activate PPARα However, GW6471, a PPARα antagonist (24), did not prevent fenofibrate from inhibiting viral infection.

To our knowledge, this is the first experimental evidence that fenofibrate can modulate RBD and ACE2 proteins and inhibit SARS-CoV-2 infection. Importantly, others have also proposed its therapeutic use in SARS-CoV-2. Fenofibrate increases the levels of the glycosphingolipid sulfatide and this has been proposed to reduce SARS-CoV-2 infection (25). SARS-CoV-2 infection is associated with overproduction of cytokines, such as TNF-α, IFN-γ, IL-1, IL-2 and IL-6, and subsequently a cytokine storm that induces several extrapulmonary complications including myocardial injury, myocarditis, acute kidney injury, impaired ion transport, acute liver injury, and gastrointestinal manifestations such as diarrhea and vomiting (26,27). Similar to dexamethasone, fenofibrate has been shown to suppress airway inflammation and cytokine release including TNF-α, IL-1 and IFN-γ in both mouse and human studies (28-30). Fenofibrate has also been shown to have antithrombotic and antiplatelet activities (31,32) reduce fibrinogen levels and increase clot permeability thereby enhancing fibrinolysis (33). These properties may reduce or prevent hypercoagulability seen in the late stage of disease in many SARS-CoV-2 patients (34). A metanalysis has also suggested fenofibrate may be useful in the treatment of Hepatitis C infection (35). Lastly, we note a preprint from the group of Nahmias that has also suggested fenofibrate may have clinical effects against SARS-CoV-2 infection which depends on the PPARα mediated alterations in host cell metabolism (36). Based on the data in this preprint, two clinical trials have been registered using fenofibrate in SARS-CoV-2 patients requiring hospitalisation (Hospital of the University of Pennsylvania (NCT04517396), and Hebrew University of Jerusalem (NCT04661930)).

Given the currently acceleration in infection and death rates observed in several countries, especially the UK, we strongly advocate clinical trials of fenofibrate in patients with SARS-CoV-2 requiring hospitalisation. Fenofibrate has a relatively safe history of use, the most common adverse effects being abdominal pain, diarrhoea, flatulence, nausea and vomiting. The half-life of fenofibric acid is 20 hours (37), allowing convenient once daily dosing. The recommended doses in the UK (up to 267 mg) provide plasma concentrations (C_max_ 70 μM, C_ss_ 50 μM) comparable to those at which we and others have seen anti-viral activity, Finally, if proven effective, fenofibrate is available as a “generic” drug and consequently is relatively cheap, making it accessible for use in all clinical settings, especially those in low and middle income countries. Further studies to clarify the precise mechanism of the anti-viral activity of fenofibrate are desirable, but this should not delay the urgent clinical evaluation of the drug to counter the current pandemic.

## ACKNOWLEDGEMENTS

Keele University funded the work performed by AR. University of Birmingham internal funds were used for work performed in Birmingham (FK SPD, ZH and HJH). CJM, MAL and MAS were funded by the BBSRC (BB/LO23717/1; BIV-HVB-2020/07/SKIDMORE and BB/S009787/1). EV and IP were funded by Bando COVID-2020-12371617, Italian Ministry of Health. Z.Y. acknowledges the Danish National Research Foundation (DNRF107) and the Lundbeck Foundation, I.B was funded by GlycoSkin H2020-ERC GAP-772735, R.K. was funded by the European Commission (GlycoImaging H2020-MSCA-ITN-721297), Y-H.C was funded by. the Innovation Fund Denmark and J.E.T. was funded by the University of Liverpool.

## CONTRIBUTIONS

AR conceived the dimerization project and performed all experimental work with the nanobit assay, immunoprecipitation studies and live whole cell binding assays (Fig 1, 2D, S2, S3, S5). FK designed the drug library and led the viral infection experiments and analysis in Birmingham (Figs 3-5, S6-S10) and performed them with SPD, ZH and HJH. MS led and ML, CJM and SG performed the biochemical studies (Fig 2A-C S4, S5B) EV led and IP performed the viral infection assays in Milan (Fig 6) Z.Y. & J.E.T. conceived, and purified the RBD-Fc protein, Y-H.C, R.K. and I.B. characterized the purified protein. AR and FK co-wrote the paper and all authors approved the final manuscript.

## Abbreviations

C_ss_: steady-state plasma concentration
C_max_: maximum plasma concentration
LgBIT: Large binary interaction technology
HiBIT: High affinity binary interaction technology
RBD: Receptor binding domain
ACE2: Angiotensin converting enzyme 2
SARS: Severe acute respiratory syndrome
ELISA: Enzyme-linked immunosorbent assay

