## Supplementary material for "The hyperlipidaemic drug fenofibrate significantly reduces infection by SARS-CoV-2 in cell culture models": supplmentary data (from another version of this preprint: version 1, 2021-01-11)

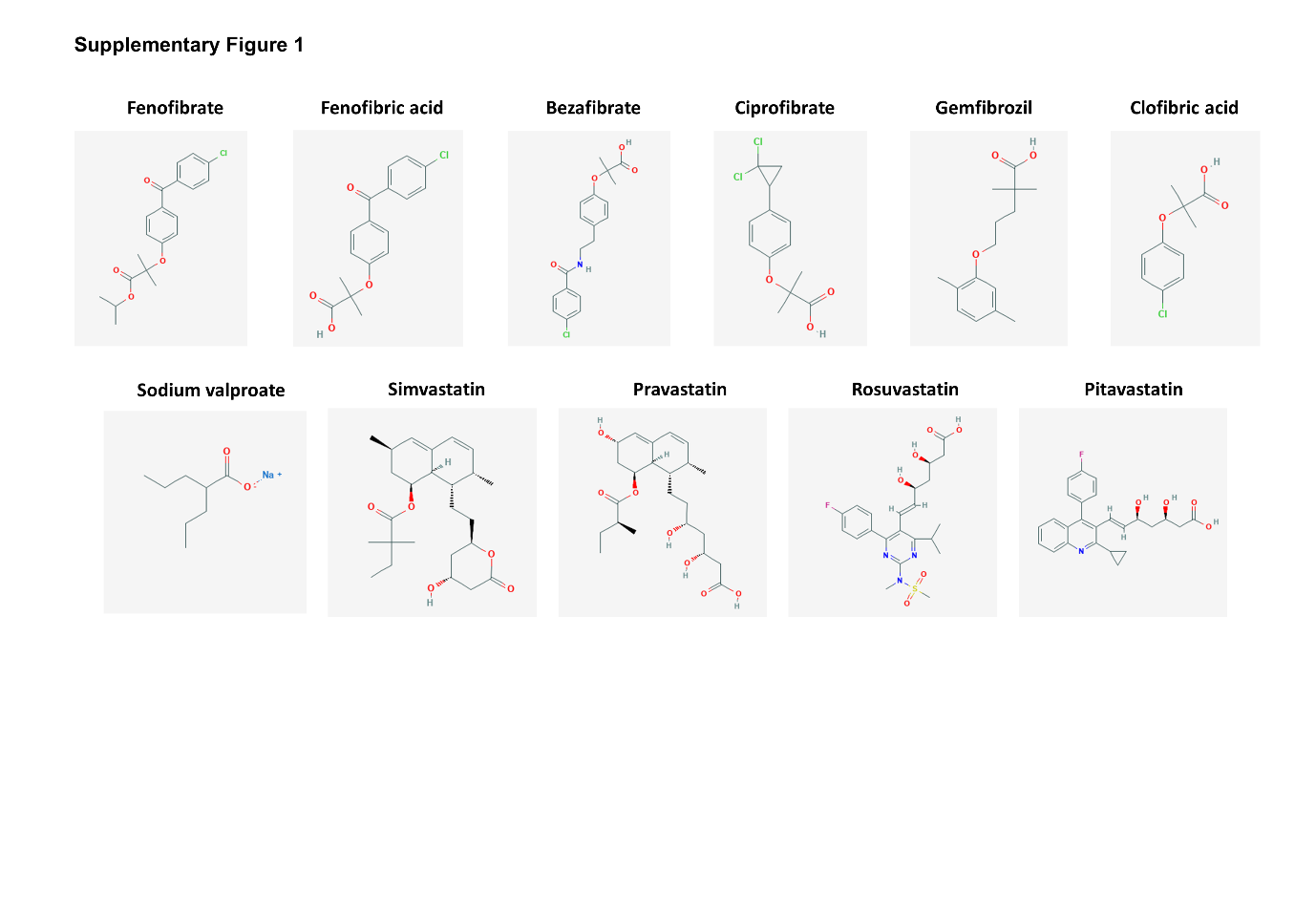

**Figure S1 Chemical structure of drugs evaluated. Structures were downloaded from pubchem.**

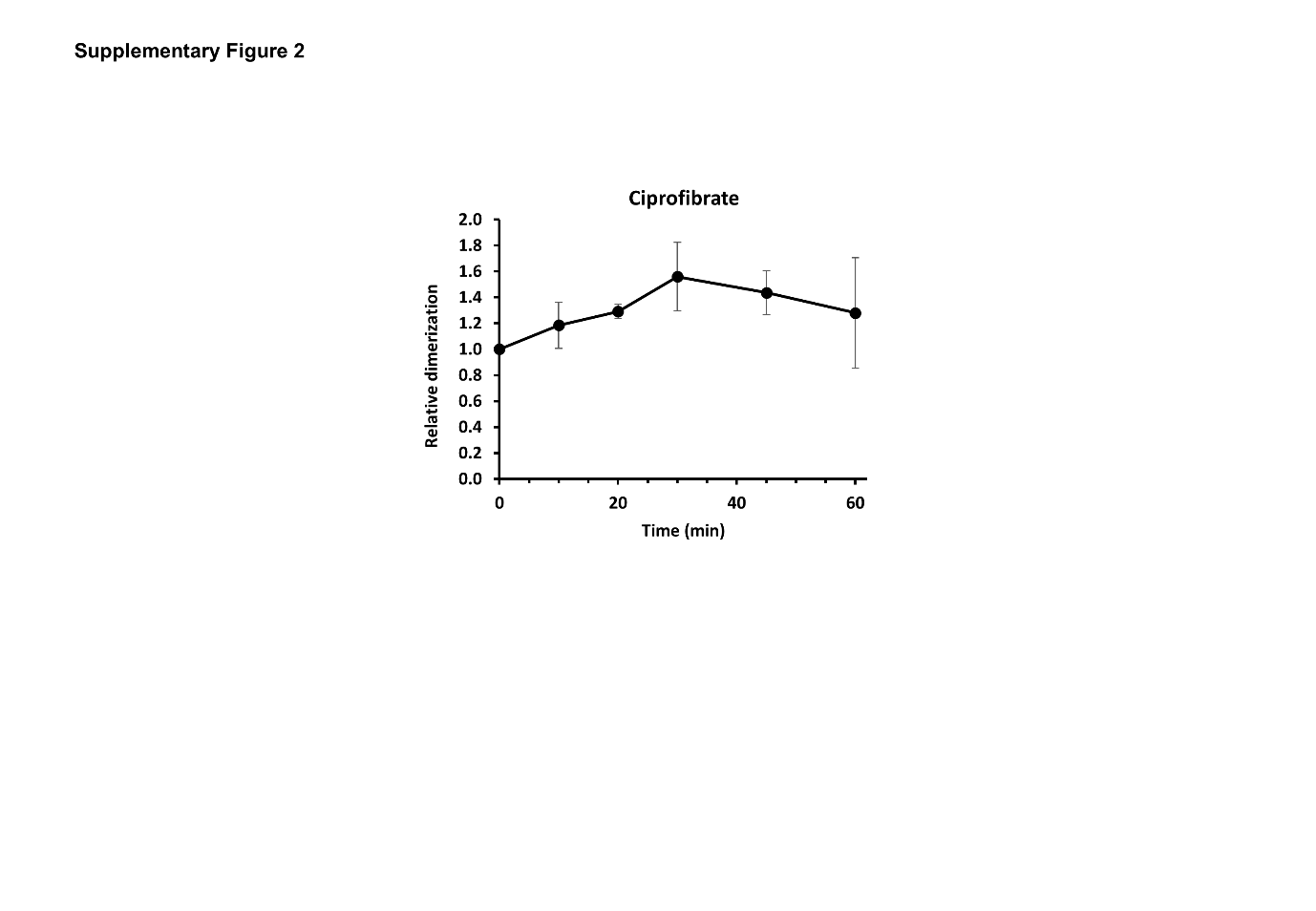
**Figure S2 Kinetics of ciprofibrate induced ACE2 dimerization.** HEK-293 cells were transfected with plasmids encoding ACE2 nanoBIT reporters. After 48 hours, the cells were treated with 230 µM ciprofibrate and the luminescence measured. The results are expressed as a proportion of that measured in cells treated with solvent (mean ± S.D>, n =3).

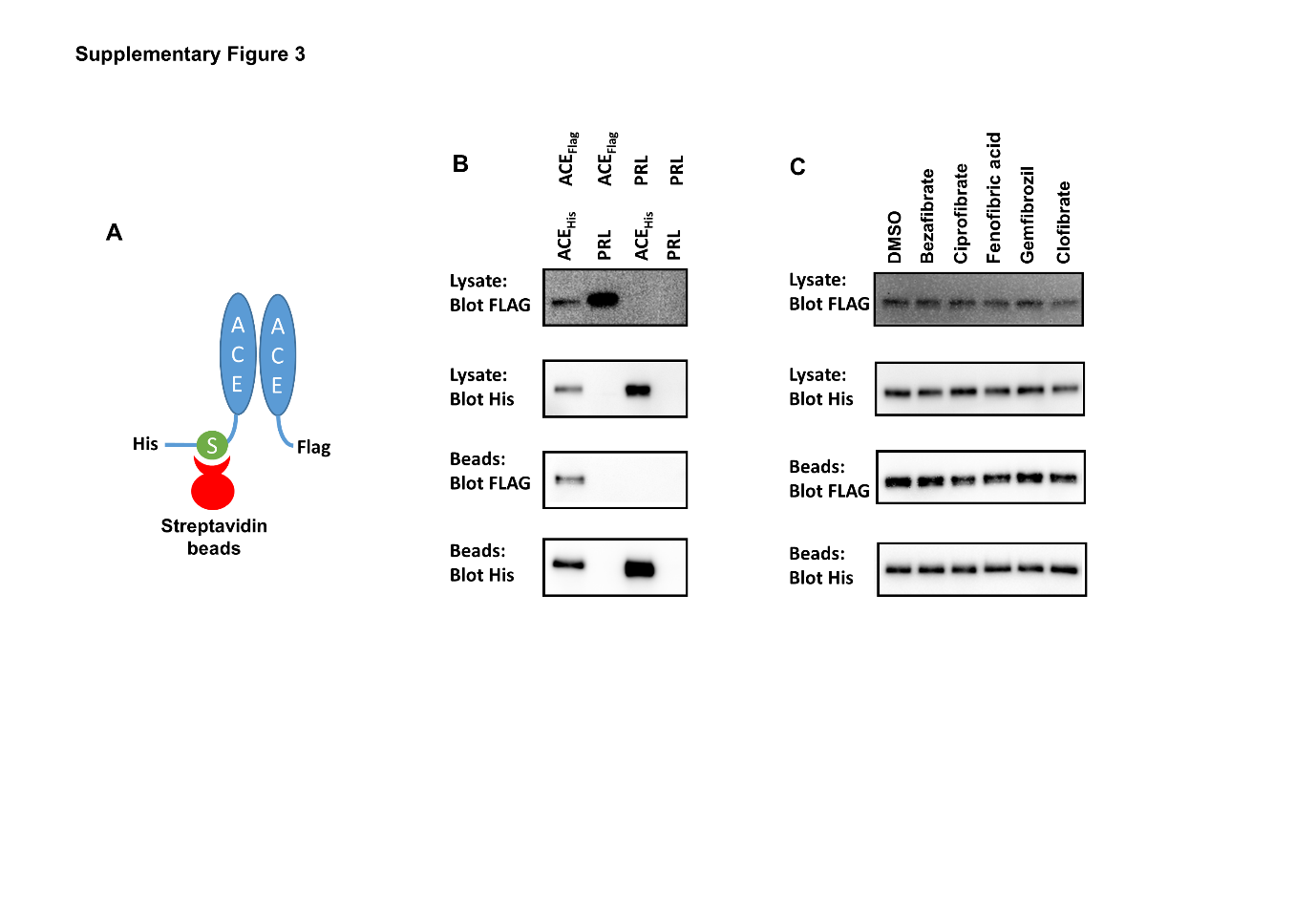

**Figure S3. Measurement of ACE2 dimers by collection on streptavidin beads. A.** Schematic showing the principle of the assay. ACE2 complexes were measured in HEK-293 cells expressing ACE2 fused to Streptavidin binding protein (S) and a His tag or encoding ACE2 with a flag tag. **B** Cells were transfected with plasmids encoding the His tagged protein (ACE_His_) or the Flag-tagged protein (ACE_Flag_) or prolactin (PRL) if one of the constructs encoding aa tagged ACE was omitted. ACE2 complexes were purified fusing streptavidin beads and analysed by immunoblotting. The results are representative of three experiments. C. HEK-293 cells expressing the tagged ACE2 proteins were exposed to the indicated drugs (230 µM) for 1 hour and the ACE2 complexes purified and analysed as described in B. The results are representative of three experiments.

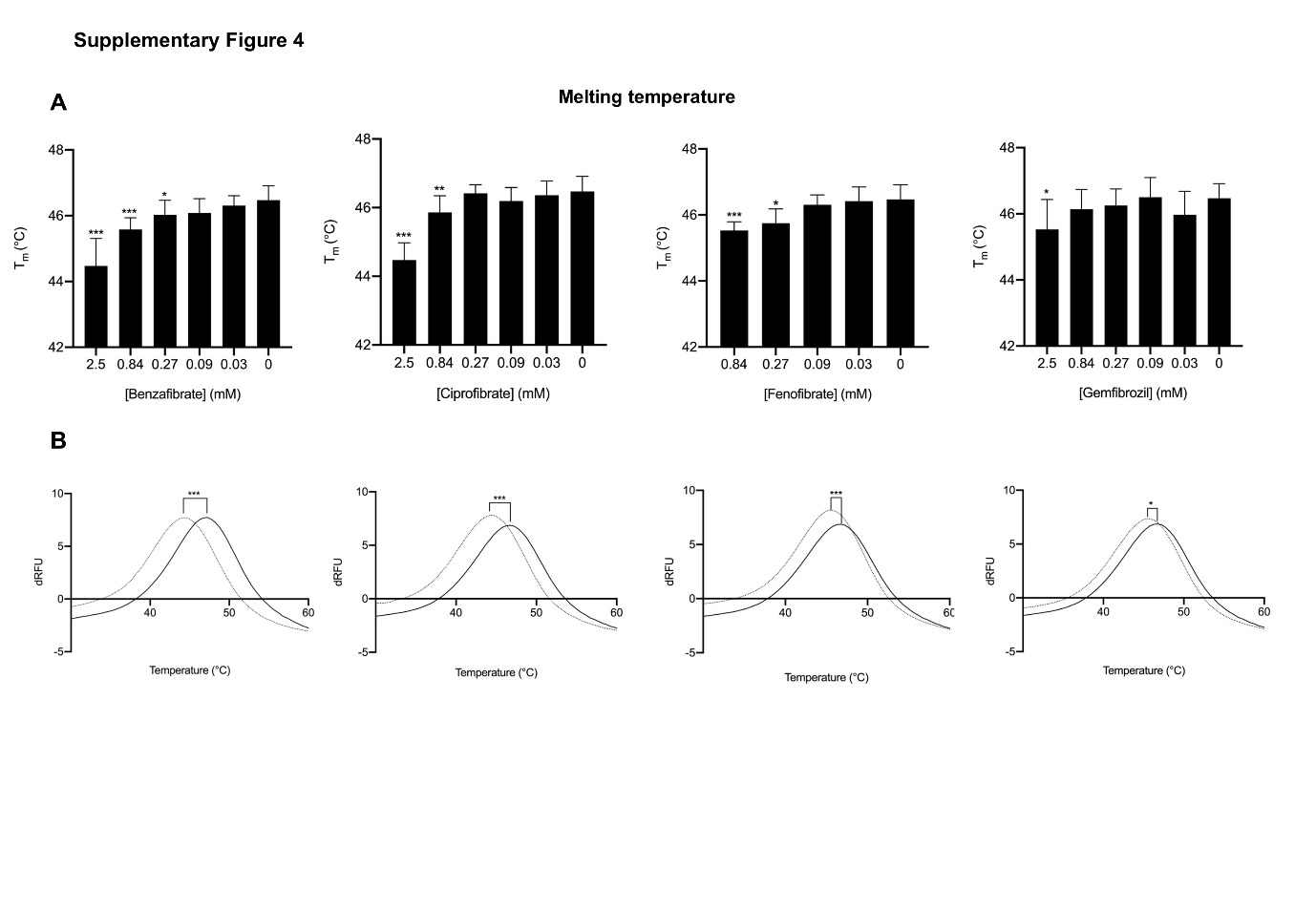

**Figure S4. Differential scanning fluorimetry.** **A**. The T_m_ of 1 μg RBD alone or with increasing concentrations of the indicated fibrate. The results (mean ± S.D., n=3) were significantly different from RBD where shown ( ***, P<0.001; **, P<0.01; *, P<0.05; paired t-test). **B.** First differential of the thermal stability of 1 μg RBD alone (solid line) or with 2.5 mM fibrate (dotted line). Fenofibrate displayed a direct interaction with SYPRO^TM^ Orange dye at the highest concentration tested (2.5 mM); as a consequence, this data point has been omitted. No interaction was observed between fenofibrate and SYPRO^TM^ Orange at concentrations ≤ 0.84 mM, at which concentration a significant difference can be overserved between the T_m_ of RBD alone and that in the presence of 0.84 mM fenofibrate, displaying a ΔT_m_ of 0.9 °C. All fibrates screened induced a comparable shift in T_m_ at 0.84 mM, with the exception of gemfibrozil, which did not display a significant change in T_m._

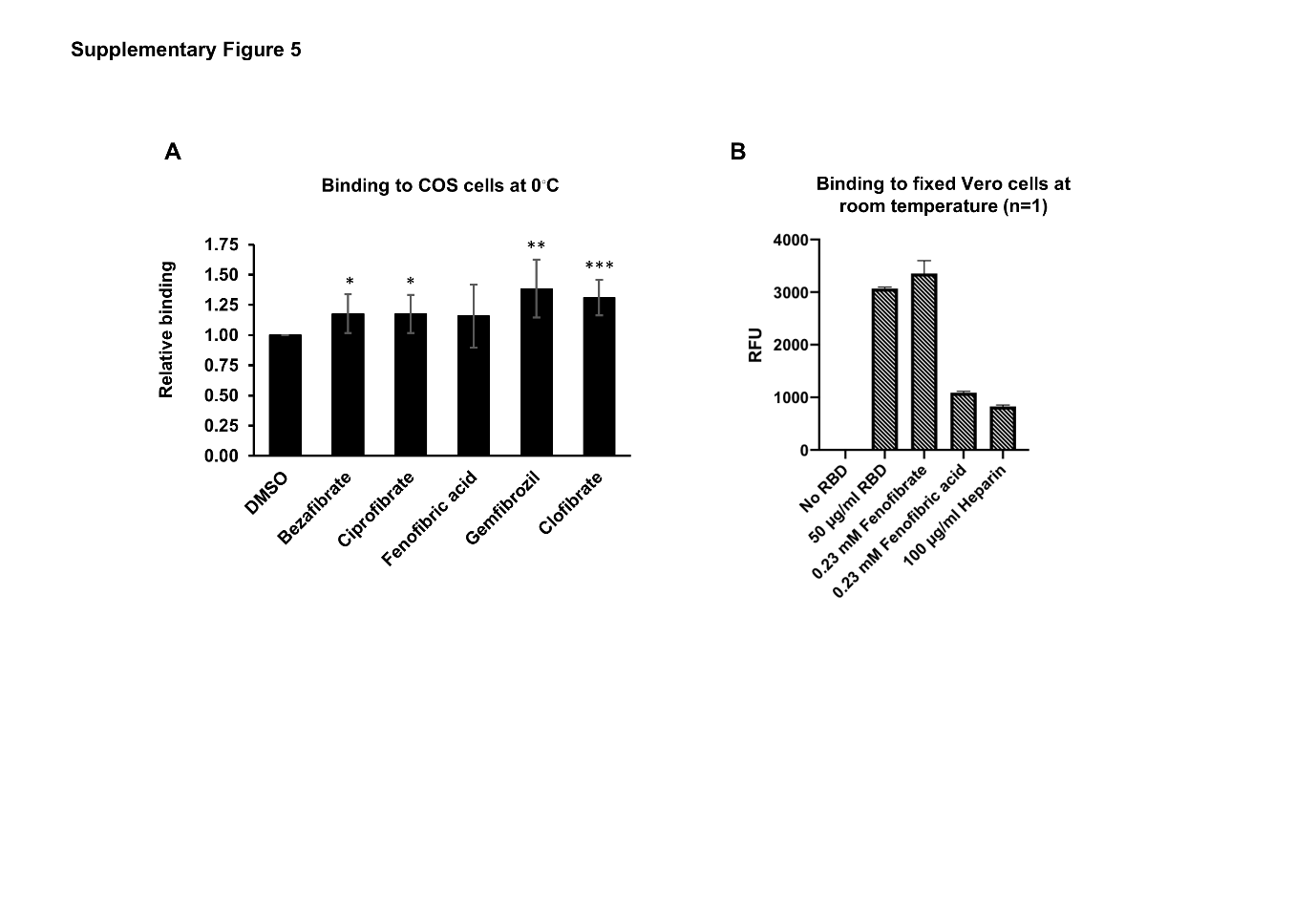

**Figure S5 Binding of RBD to ACE2. A.** Binding to live COS cells at 0°C. COS cells were transfected with ACE2 and binding of RBD-HiBIT, separately expressed in COS cells, measured as described(12). The results (mean ± S.D, n=7) are expressed as proportion of the binding to cells treated with solvent alone and are different to it where shown (*, P<0.05; **, P<0.01, ***. P<0.005). **B.** Binding or RBD to ACE2 in fixed Vero cells was measured in the presence of the indicated drug. This is a preliminary experiment performed once.

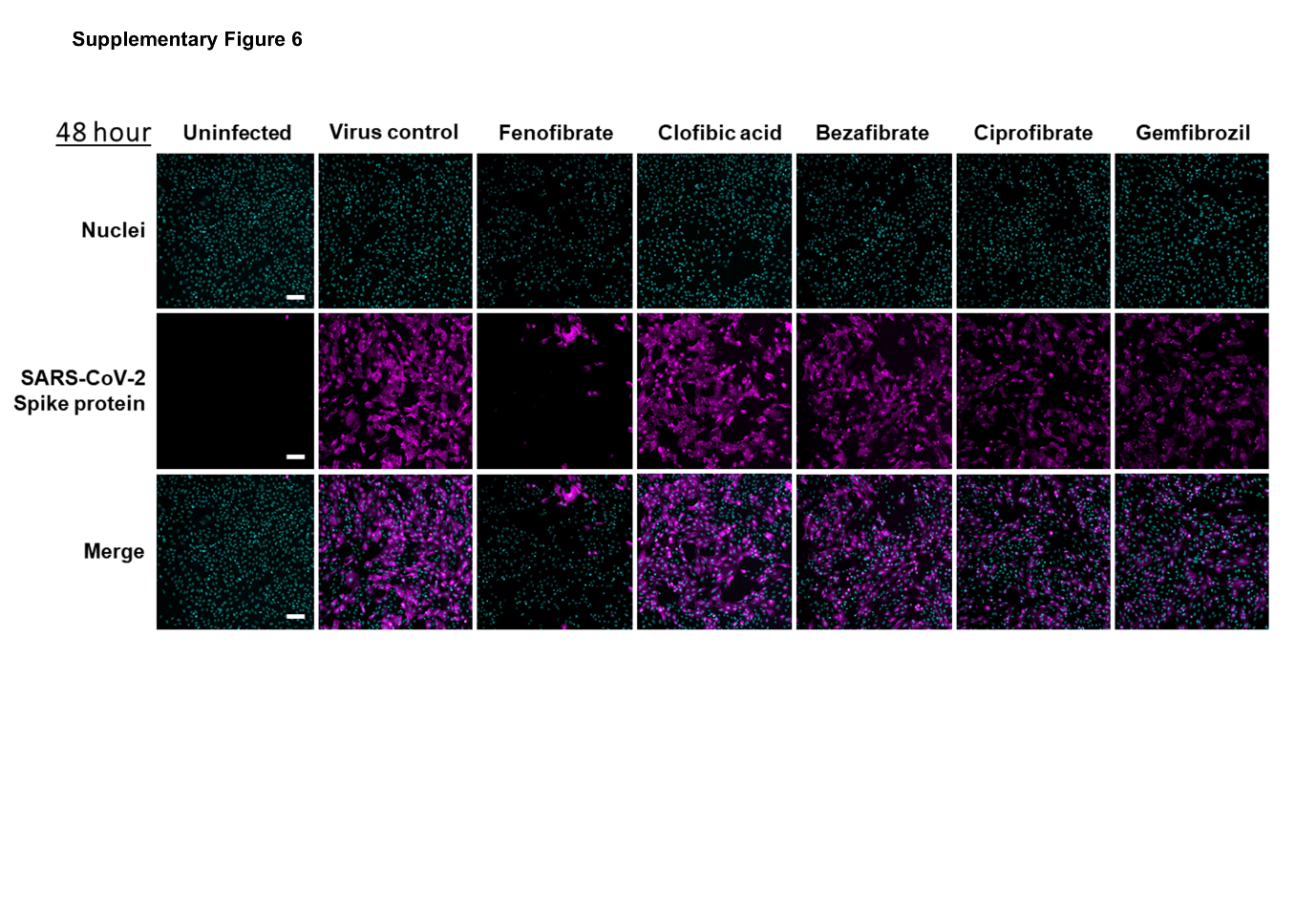

**Figure S6. Fenofibrate reduces SARS-CoV-2 Vero infection in Vero cells as measured by Spike protein staining**

Vero cells were plated in 96 well plates at 8x103 cells/well for 24 hours before inoculating with SARS-CoV-2 strain hCOV-19/England/2/2020 at 167 IU/well with 230 µM fibrates or DMSO equivalent in a total volume of 50ml. After 48 hours, supernatant was removed and cells were fixed in ice-cold methanol. Cells were then stained with rabbit anti-SARS-COV-2 spike protein, subunit 1 followed by Alexa Fluor 555-conjugated goat anti-rabbit IgG secondary antibody. Cell nuclei were visualised with Hoechst 33342. Cells were washed with PBS and then imaged and analysed using a Thermo Scientific CellInsight CX5 High-Content Screening (HCS) platform. Infected cells were scored by perinuclear fluorescence above a set threshold determined by positive (untreated) and negative (uninfected) controls. A minimum of 9 fields and 5000 nuclei per well in triplicate or quadruplicate wells per treatment were scored in each experiment. All experiments were performed n=2-4 with 3-4 repeats in each experiment.

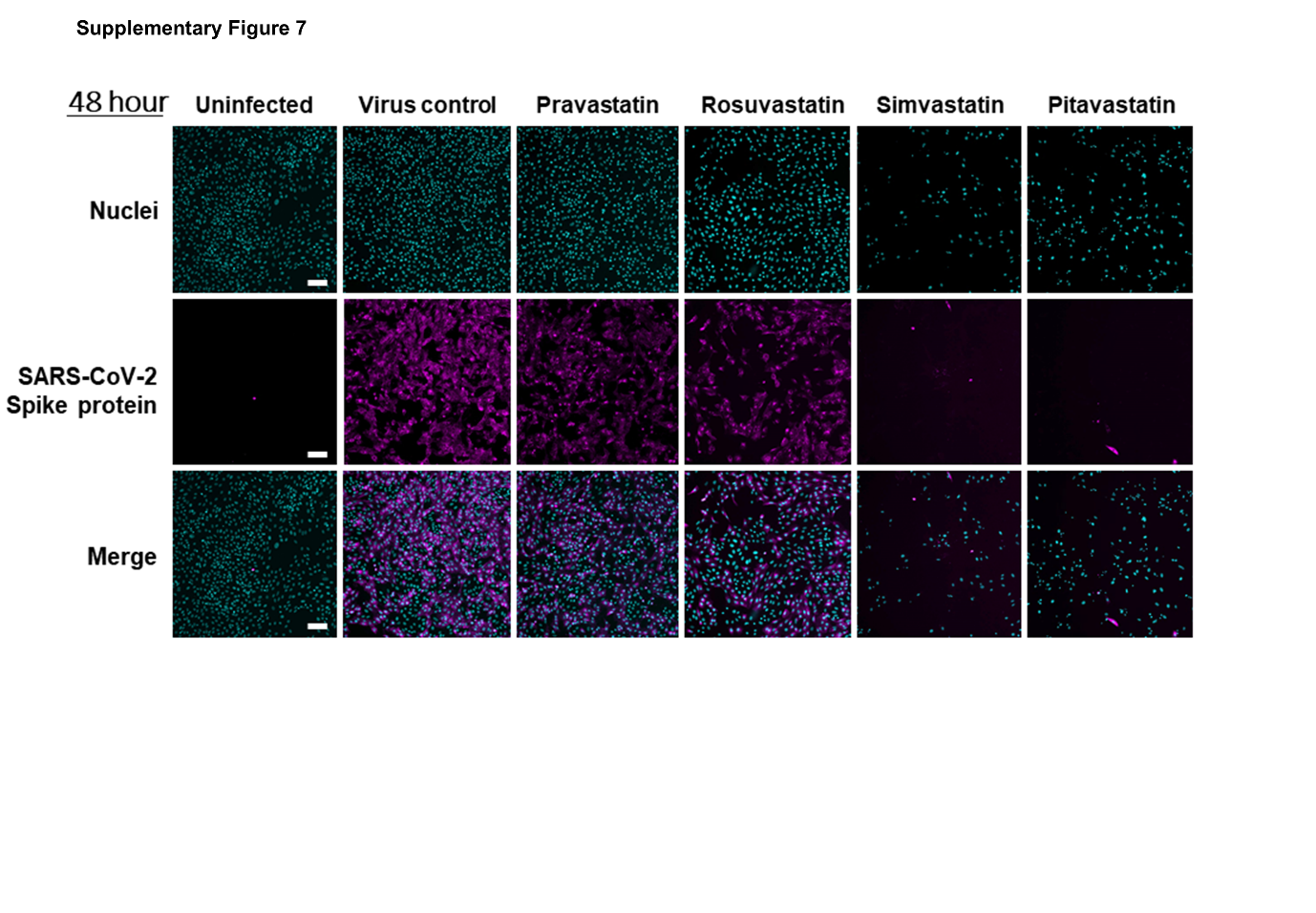

**Figure S7. Statins do not reduce infection of Vero cells by SARS-CoV-2 Vero**

Vero cells were plated in 96 well plates at 8x10^3^ cells/well for 24 hours before inoculating with SARS-CoV-2 strain hCOV-19/England/2/2020 at 167 IU/well with 100nM statins or DMSO equivalent in a total volume of 50ml. After 48 hours, supernatant was removed and cells were fixed in ice-cold methanol. Cells were then stained with rabbit anti-SARS-COV-2 spike protein, subunit 1 followed by Alexa Fluor 555-conjugated goat anti-rabbit IgG secondary antibody. Cell nuclei were visualised with Hoechst 33342. Cells were washed with PBS and then imaged and analysed using a Thermo Scientific CellInsight CX5 High-Content Screening (HCS) platform. Infected cells were scored by perinuclear fluorescence above a set threshold determined by positive (untreated) and negative (uninfected) controls. A minimum of 9 fields and 5000 nuclei per well in triplicate or quadruplicate wells per treatment were scored in each experiment. All experiments were n=2-4 with 3-4 repeats in each experiment.

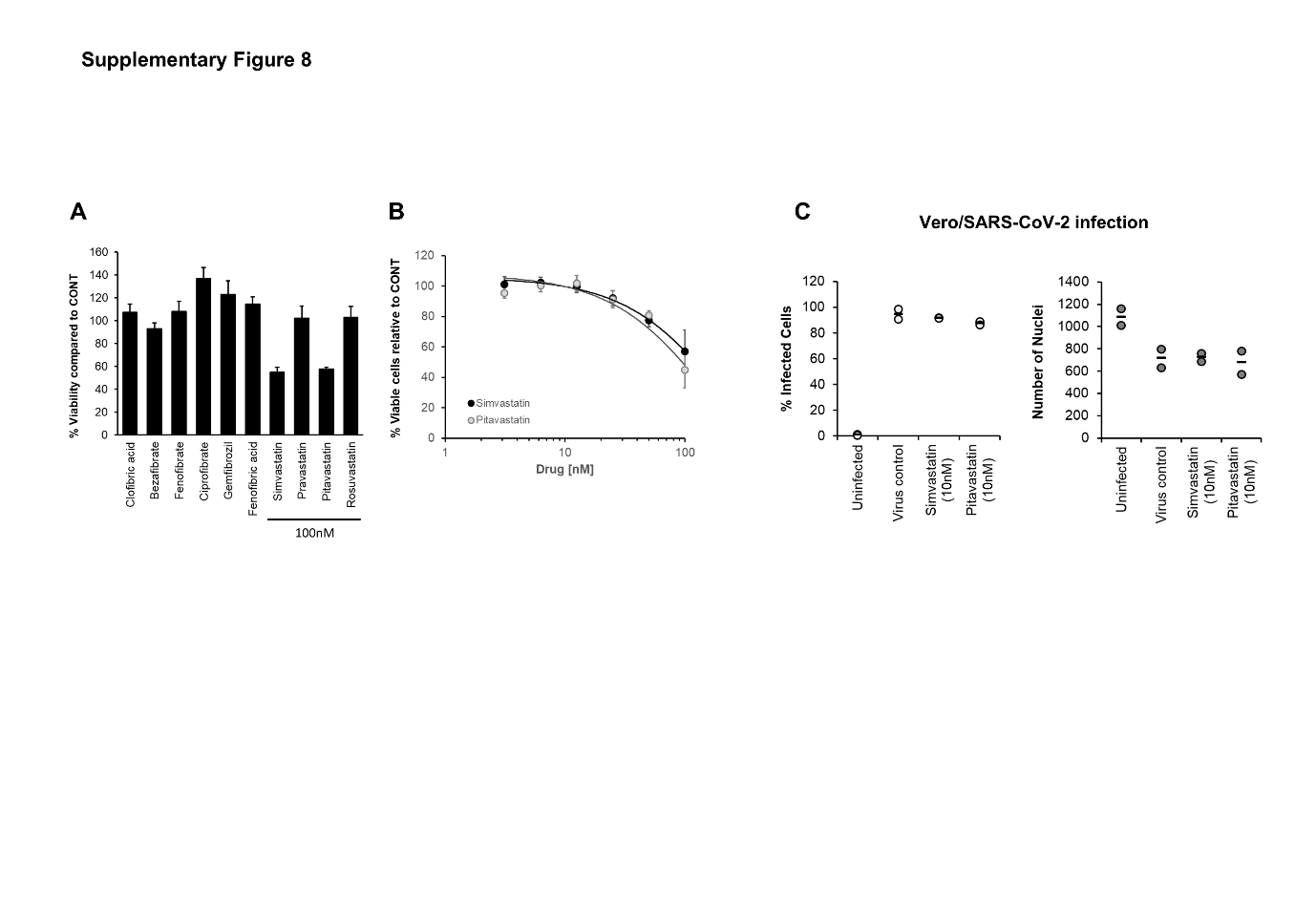

**Figure S8. Statins do not reduce infection of Vero cells with SARS-CoV-2**

Vero cells were plated in 96 well plates at 8x10^3^ cells/well for 24 hours before treating with either 100nM statins (A) or doubling dilutions of simvastatin or pitavastatin (B) in 200ul. Viability was assessed at 48 hours using Cell-titre Blue (Promega) assay and viability calculated relative to DMSO controls. (C) Vero cells were infected with SARS-CoV-2 strain hCOV-19/England/2/2020 at 167 IU/well with 10nM statins in a total volume of 50ml. After 48 hours, supernatant was removed and cells were fixed in ice-cold methanol. Cells were then stained with rabbit anti-SARS-COV-2 spike protein, subunit 1 followed by Alexa Fluor 555-conjugated goat anti-rabbit IgG secondary antibody. Cell nuclei were visualised with Hoechst 33342. Cells were washed with PBS and then imaged and analysed using a Thermo Scientific CellInsight CX5 High-Content Screening (HCS) platform. Infected cells were scored by perinuclear fluorescence above a set threshold determined by positive (untreated) and negative (uninfected) controls. A minimum of 9 fields and 5000 nuclei per well in triplicate or quadruplicate wells per treatment were scored in each experiment. All experiments were performed n=2-4 with 3-4 repeats in each experiment.

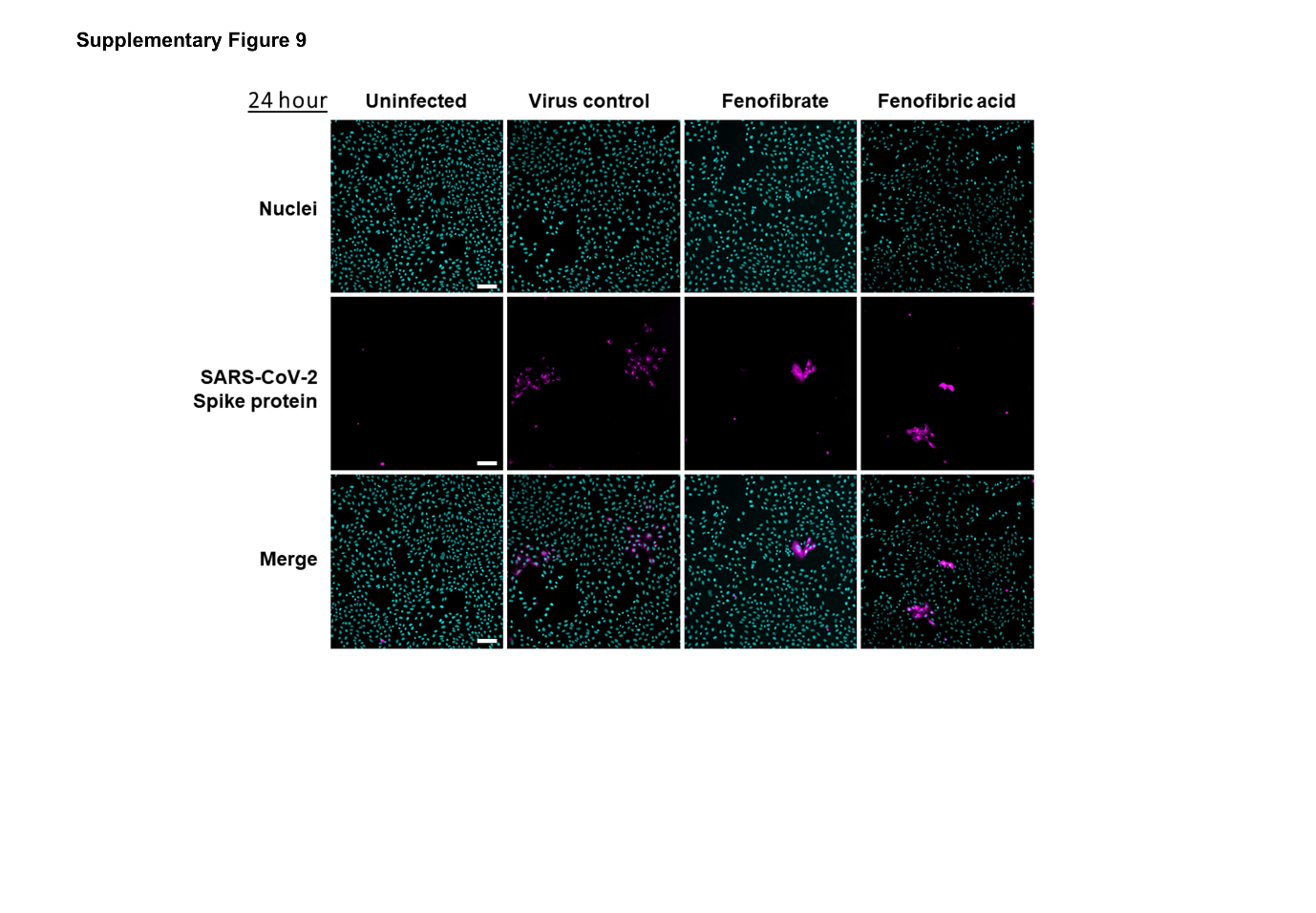
**Figure S9. Fenofibrate reduces infection rates of Vero cells by SARS-CoV-2 within 24 hours**

Vero cells were plated in 96 well plates at 8x10^3^ cells/well for 24 hours before inoculating with SARS-CoV-2 strain hCOV-19/England/2/2020 at 167 IU/well with 100nM statins or DMSO equivalent in a total volume of 50ml. After 24 hours, supernatant was removed and cells were fixed in ice-cold methanol. Cells were then stained with rabbit anti-SARS-COV-2 spike protein, subunit 1 followed by Alexa Fluor 555-conjugated goat anti-rabbit IgG secondary antibody. Cell nuclei were visualised with Hoechst 33342. Cells were washed with PBS and then imaged and analysed using a Thermo Scientific CellInsight CX5 High-Content Screening (HCS) platform. Infected cells were scored by perinuclear fluorescence above a set threshold determined by positive (untreated) and negative (uninfected) controls. A minimum of 9 fields and 5000 nuclei per well in triplicate or quadruplicate wells per treatment were scored in each experiment. All experiments were performed n=2-4 with 3-4 repeats in each experiment.

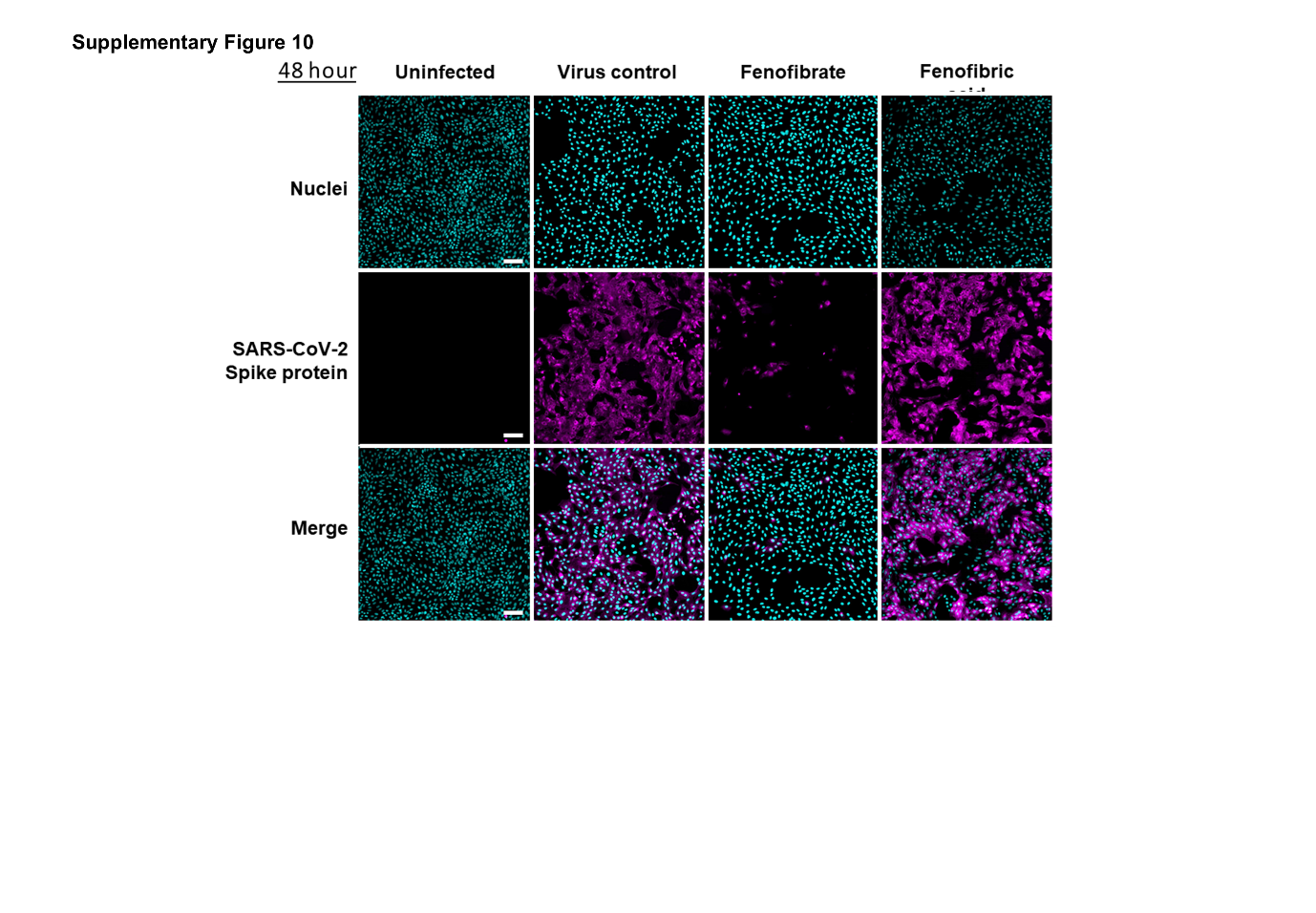

**Figure S10. Fenofibrate reduces SARS-CoV-2 infections in Vero cells at 48 hours**

Vero cells were plated in 96 well plates at 8x10^3^ cells/well for 24 hours before inoculating with SARS-CoV-2 strain hCOV-19/England/2/2020 at 167 IU/well with 100nM statins or DMSO equivalent in a total volume of 50ml. After 48 hours, supernatant was removed and cells were fixed in ice-cold methanol. Cells were then stained with rabbit anti-SARS-COV-2 spike protein, subunit 1 followed by Alexa Fluor 555-conjugated goat anti-rabbit IgG secondary antibody. Cell nuclei were visualised with Hoechst 33342. Cells were washed with PBS and then imaged and analysed using a Thermo Scientific CellInsight CX5 High-Content Screening (HCS) platform. Infected cells were scored by perinuclear fluorescence above a set threshold determined by positive (untreated) and negative (uninfected) controls. A minimum of 9 fields and 5000 nuclei per well in triplicate or quadruplicate wells per treatment were scored in each experiment. All experiments were performed n>3 times with 3-4 repeats in each experiment.
